# Neurogliaform cells mediate interhemispheric modulation of sensory-evoked activity in cortical pyramidal neurons

**DOI:** 10.1101/2025.04.30.651595

**Authors:** Foivos Markopoulos, Ronan Chéreau, Federico Brandalise, Vaibhav Chippalkatti, Julien Prados, Alexandre Dayer, Anthony Holtmaat

**Affiliations:** Department of Basic Neurosciences, and Neurocenter, Faculty of Medicine, University of Geneva, Geneva, Switzerland; Bioinformatics support platform, Faculty of Medicine, University of Geneva, Geneva, Switzerland; Department of Psychiatry, University of Geneva, Geneva, Switzerland

## Abstract

Integration of bilateral sensory inputs requires effective communication between brain hemispheres. This interhemispheric communication is essential for sensory perception and involves reciprocal connections between homotopic sensory areas. A key role in this process is attributed to interhemispheric inhibition which, owing to its long-lasting form, is posited to operate largely through neurogliaform cells (NGCs). However, direct evidence on the role of NGCs in interhemispheric inhibition is missing. Here we show that NGCs in the mouse barrel cortex (BC) are engaged by interhemispheric callosal projections to modulate pyramidal neuron (PN) activity and sensory perception. Using optogenetics, *ex vivo* whole-cell recordings, and *in vivo* calcium imaging, we found that layer 1 and layer 2/3 (L1-3) NGCs are strongly activated by the callosal and suppressed by the thalamocortical pathway, suggesting that NGCs encode ipsilateral rather than contralateral whisker stimuli. We also found that direct stimulation of L1-3 NGCs modulates whisker-evoked activity in L2/3 and L5b PNs, and increases the perceptual threshold in a whisker-deflection detection task. Furthermore, these effects were recapitulated by direct stimulation of callosal projections and deflection of the ipsilateral whiskers respectively, suggesting that the effect of the callosal pathway on sensory perception is mediated by NGCs. Our results not only prove that NGCs mediate interhemispheric inhibition, but also demonstrate their role in sensory perception via modulation of the main units involved in cortical input and output.

## INTRODUCTION

Sensory stimuli appearing at either side of the mammalian body are processed in both the contralateral and ipsilateral cerebral hemisphere. It is thought that this facilitates the uniform perception of bilateral sensory inputs, especially those close to the body’s midline ^1^. Thus, neurons in sensory cortices of each hemisphere have receptive fields that span the contra and ipsilateral sides of or hemifields around the body ^2,3^. The bilateral cortical processing and integration of sensory input relies strongly on interhemispheric communication via transcallosal fibers, which can be both excitatory and inhibitory in nature ^4–8^. Even though callosal afferents span all cortical layers, interhemispheric inhibition was shown to strongly implicate layer 1 interneurons (L1 INs) ^7^. A large fraction of L1 INs in the hindlimb area of the primary somatosensory cortex (S1HL) were shown to receive monosynaptic inputs from homotopic callosal afferents, and accordingly, ∼40% of L1 INs in S1 showed a response to ipsilateral sensory stimulation ^7^. This study also showed that ipsilateral sensory stimulation appears to elicit long-lasting inhibition and suppresses contralateral sensory-evoked activity of L5 PN dendrites in S1HL, in a GABA_B_R-dependent manner ^7^. Therefore, the implicated L1 INs were postulated to be neurogliaform cells (NGCs) ^7^, which is an inhibitory cell type that is characterized by their tendency to evoke slow inhibition through GABA_B_ receptors on their post-synaptic targets ^9^. However, the NGC-selective population responses in the context of interhemispheric cortical processing are lacking, as specific labeling or interrogation of these interneurons has been hindered by the lack of transgenic mouse lines that would provide selective genetic access. In the absence of such tools, and since NGCs are more abundant in L1, several studies have used L1 INs, which have recently become tractable through the NDNF-Cre mouse line ^10^, as a proxy for NGCs ^11,12^. Using these tools, it has been shown that in the primary auditory and visual cortices (A1 and V1), L1 NDNF INs receive monosynaptic inputs from thalamocortical afferents ^13,14^ and are activated upon sensory stimulation ^10,11^. Recent work in which genetically identified L1 NDNF INs were interrogated for their role in bilateral sensory processing corroborate the former finding that they play a role in interhemispheric inhibition ^12^. However, the latter study suggests that interhemispheric interactions suppress L2/3 PN activity, whereas the former study deemed the inhibition selective for L5 PNs and concluded that L2/3 PNs must receive a dominant level of excitation from transcallosal inputs ^15^. Therefore, it remains unclear whether previous findings hold true specifically for L1 NGCs which correspond to only ∼30% of L1 NDNF or ∼40% of all L1 INs ^16^. Moreover, since NGCs span all cortical layers ^17^, the input connectivity and sensory responses of deeper layer NGCs remain unknown.

Here, we set out experiments to more precisely investigate the synaptic connectivity between transcallosal inputs and NGCs in the mouse S1 barrel cortex, and map their role in bilateral sensory input integration in awake mice. To this end, we use a transgenic mouse line approach that allows selective labeling of NGCs ^17,18^, and which we recently used for imaging NGC activity *in vivo* ^19^. We report data on the interhemispheric and thalamic control of L1-3 NGC activity, *ex vivo* and *in vivo*, and unravel the modulatory effect of L1-3 NGCs on sensory-evoked activity of L2/3 and L5b neurons. We provide evidence that this effect is recapitulated by sensory activation of the callosal pathway.

## RESULTS

### Transcallosal slow inhibition of cortical pyramidal neurons is mimicked by activation of NGCs

To investigate whether NGCs mediate transcallosal long-lasting inhibition, we first sought to test whether the previously documented transcallosal inhibition of L5 pyramidal dendrites also applies to L2/3 pyramidal neurons in the S1 BC, and how this compares to the synaptic drive they receive from thalamocortical (TC) input. To this end, the light-gated ion channel channelrhodopsin-2 (ChR2) ^20^ was unilaterally expressed using targeted injections of recombinant adeno-associated viral vectors (AAV). Vectors were injected either in the somatosensory thalamus (ventroposterior [VPM] and posteromedial [POm]) of the same hemisphere as the subsequent recording site, or in the S1 BC of the other hemisphere (Figure 1A and B). We recorded intracellular responses from L2/3 PNs (Figure 1A-C) in brain slices upon optical stimulation of the ChR2-expressing TC afferents or the callosal BC afferents (BCc) (Figure 1A-B). L2/3 PNs were identified based on their typical shape and firing pattern. In current clamp recordings, upon continuous current injection to maintain membrane potentials at ∼-55 mV, optical stimulation of TC afferents evoked robust excitatory postsynaptic potentials (EPSPs) with short and long-lasting components (similar to sensory-evoked responses *in vivo* ^21^, but not inhibitory postsynaptic potentials (IPSPs; Figure 1D). In contrast, optical stimulation of BCc afferents evoked robust EPSPs followed by robust IPSPs (Figure 1E). Bath application of the GABA_B_R antagonist CGP-55845 reduced the late component of the IPSPs by ∼40% (Figure 1E).

**Figure 1.**
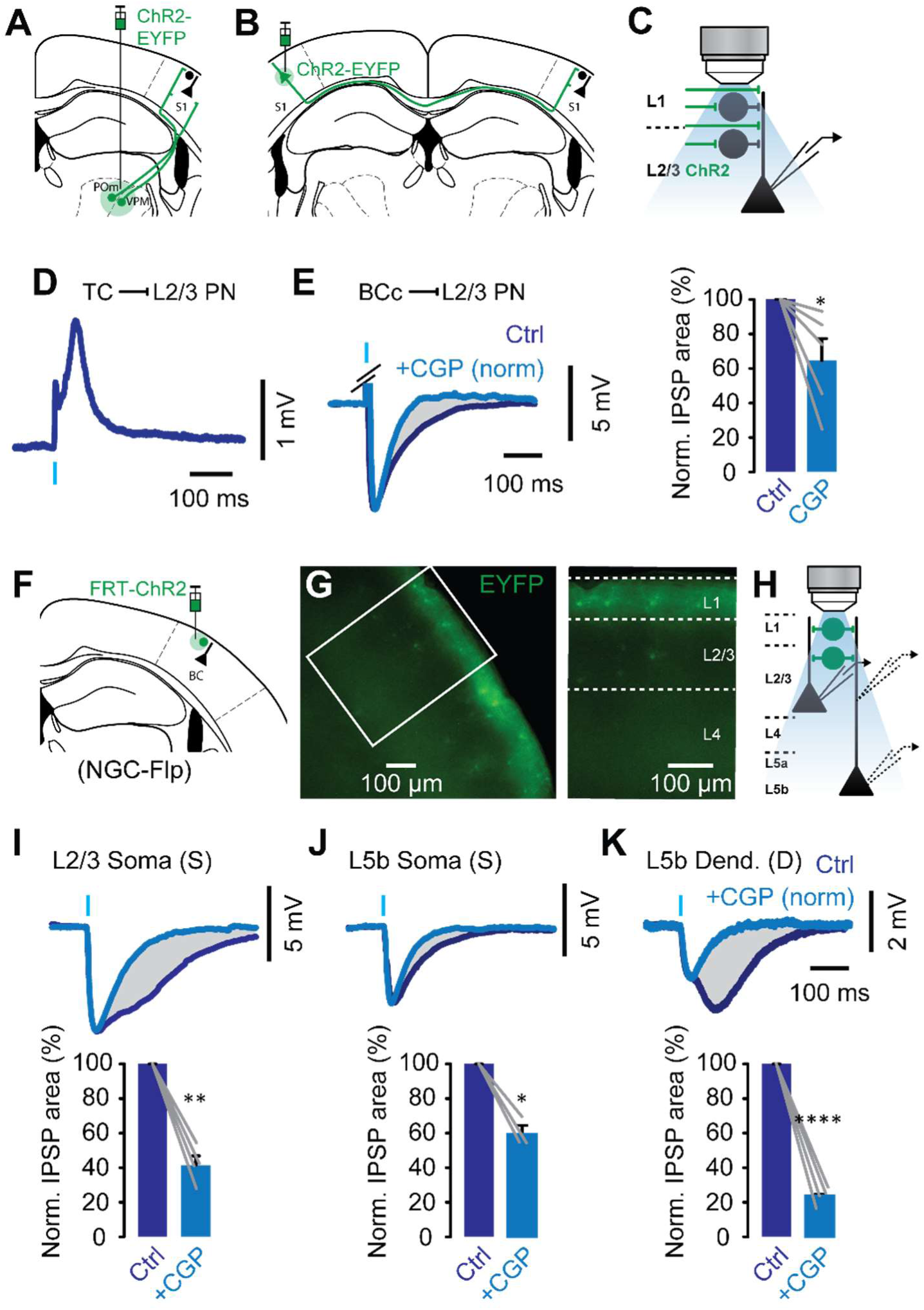
NGC-mediated slow inhibition in pyramidal neurons. (A and B) Schematic representation of AAV-directed ChR2 expression in TC and BCc afferents, in WT mice. (C) Schematic representation of whole-cell current clamp recordings from L2/3 PN somata upon optical stimulation of TC and BCc afferents. (D) Representative example of average EPSPs recorded at Vm = ∼55 mV upon optical stimulation of TC afferents. (E) Left, representative example of average IPSPs recorded at Vm = ∼55 mV upon optical stimulation of BCc afferents before (dark blue) and after bath application of CGP-55845 (1 µM; light blue). Preceding EPSPs were covered for clarity. Grayed area indicates the abolished GABA_B_R-dependent component. Right, normalized areas of average IPSPs before and after CGP-55845 (*n = 5* cells from *N = 5* mice). *P = 0.0499, paired t-test. (F) Schematic representation of AAV-directed ChR2 expression in L1-3 NGCs, in NGC-Flp mice. (G) Epifluorescence images of ChR2-expressing NGCs in the BC of recorded slices. (H) Schematic representation of whole-cell current clamp recordings from L2/3 and L5b PN somata, and L5b PN trunks, upon optical stimulation of L1-3 NGCs, in separate experiments. (I, J, and K) Top, Representative examples of average IPSPs recorded at Vm = ∼55 mV, upon optical stimulation of L1-3 NGCs before (dark blue) and after bath application of CGP-55845 (1 µM; light blue). Light blue traces are normalized to the amplitude of the first IPSP of dark blue traces. Grayed areas indicate the abolished GABA_B_R-dependent components. Bottom, normalized areas of average IPSPs before and after CGP-55845 (*n = 4* cells from *N = 2* mice for I, *n = 3* cells from *N = 2* mice for J, and *n = 4* dendrites from *N = 3* mice for K). **P = 1.7 x 10^-3^ for I, *P = 0.0147 for J, ****P = 8.5 x 10^-5^ for K, paired t-test.

Together, these data show that BCc afferents can drive a long-lasting inhibition of L2/3 PNs via a GABA_B_R-dependent mechanisms, in line with the earlier notion of transcallosal L5 PN inhibition ^7^. In contrast, this form of inhibition is not apparent when TC afferents are stimulated.

We then sought to investigate whether NGCs could specifically mediate the long-lasting inhibitory response that was observed upon stimulation of BCc afferents. To selectively target NGCs, we utilized a *Hmx3*-Cre mouse line, which is a reporter line that was previously employed to track the migration and development of cortical NGCs originating from the preoptic area ^17,18^. Since *Hmx3*-Cre mice express Cre only at embryonic stages, it was crossbred with mice harboring the Cre-conditional *Rosa-26-CAG-LSL-Flp* allele to generate offspring that stably expresses the recombinase flipase (Flp) in the mature population of NGCs, which we hereafter refer to as NGC-Flp mice. ChR2 expression in L1-3 NGCs was achieved using targeted injections of a Flp-dependent AAV in the BC of these NGC-Flp mice ^19^. In successful injections, a sparse and robust ChR2-YFP expression was observed in L1-3 corresponding to NCGs (Figure 1G). In brain slices, L2/3 PNs were recorded at the soma, whereas L5b PNs were recorded at both the soma and dendrite, in separate sets of experiments (Figure 1H).

In current clamp recordings, upon continuous current injection to maintain membrane potentials at ∼-55 mV, optical stimulation of NGCs evoked robust IPSPs in all recorded sites (Figure 1I-K). These IPSPs had similar kinetics to those previously described using paired recordings between NGCs and PNs ^22^. Bath application of the GABA_B_R antagonist CGP-55845 significantly reduced their duration and revealed a GABA_B_R-dependent slow component (Figure 1I-K). In some cells we subsequently applied the GABA_A_R antagonist Gabazine (SR-95531) which completely abolished the remaining IPSPs, indicating that they were GABA_A_R-dependent (Figure S1A). However, we noticed that the proportion of GABA_B_R- and GABA_A_R-dependent components differed between recording sites (Figure S1B top). To quantify these differences, we computed the GABA_B_/GABA_A_ ratio for each recording site (Methods). Our analysis showed that this ratio was significantly higher for L2/3 PN somata and L5b PN dendrites compared to L5b PN somata, as well as for L5b PN dendrites compared to L2/3 PN somata (Figure S1B bottom). Notably, the striking difference in the GABA_B_/GABA_A_ ratio between L5b PN dendrites and somata, suggests that NGCs control these neurons almost exclusively via GABA_B_Rs in the apical dendrites (Figure 1J,K). This is in line with a recent study highlighting the role of NGCs and GABA_B_Rs in dendro-somatic synergy of L5 PNs ^23^.

This data support the notion that NGCs provide a long-lasting inhibition of L2/3 and L5 neurons, particularly onto their dendrites, that is dependent on GABA_B_R. The profile of this long-lasting inhibition bears similarities to the inhibition that is driven by BCc afferents, which suggests that NGCs are the predominant source of BCc-mediated inhibition of BC PNs.

### NGCs exhibit a higher E-I ratio upon optogenetic stimulation of callosal compared to thalamocortical afferents

Next, we sought to investigate the synaptic inputs that L1-3 NGCs receive from BCc afferents as compared to TC afferents. To address this, we expressed TdTomato (TdT) in cortical NGCs, in order to visually identify them, again by using the *Hmx3*-Cre mouse line and crossing it with the *Rosa-26*-CAG-LSL-TdT mouse line. In this offspring (termed NGC-TdT mice), ChR2 was unilaterally expressed in somatosensory thalamus, or S1 BC, using targeted injections of AAV (Figure 2A and D). Successful injections were validated by the observation of robust expression of ChR2-EYFP in neurons of the injected areas, as well as by the distinct expression patterns within the TC and BCc afferents in the BC (Figure 2B,C and E,F). We performed intracellular recordings of TdT-positive L1-3 NGCs in brain slices to measure response properties upon optical stimulation of TC and BCc afferents (Figure 2G).

**Figure 2.**
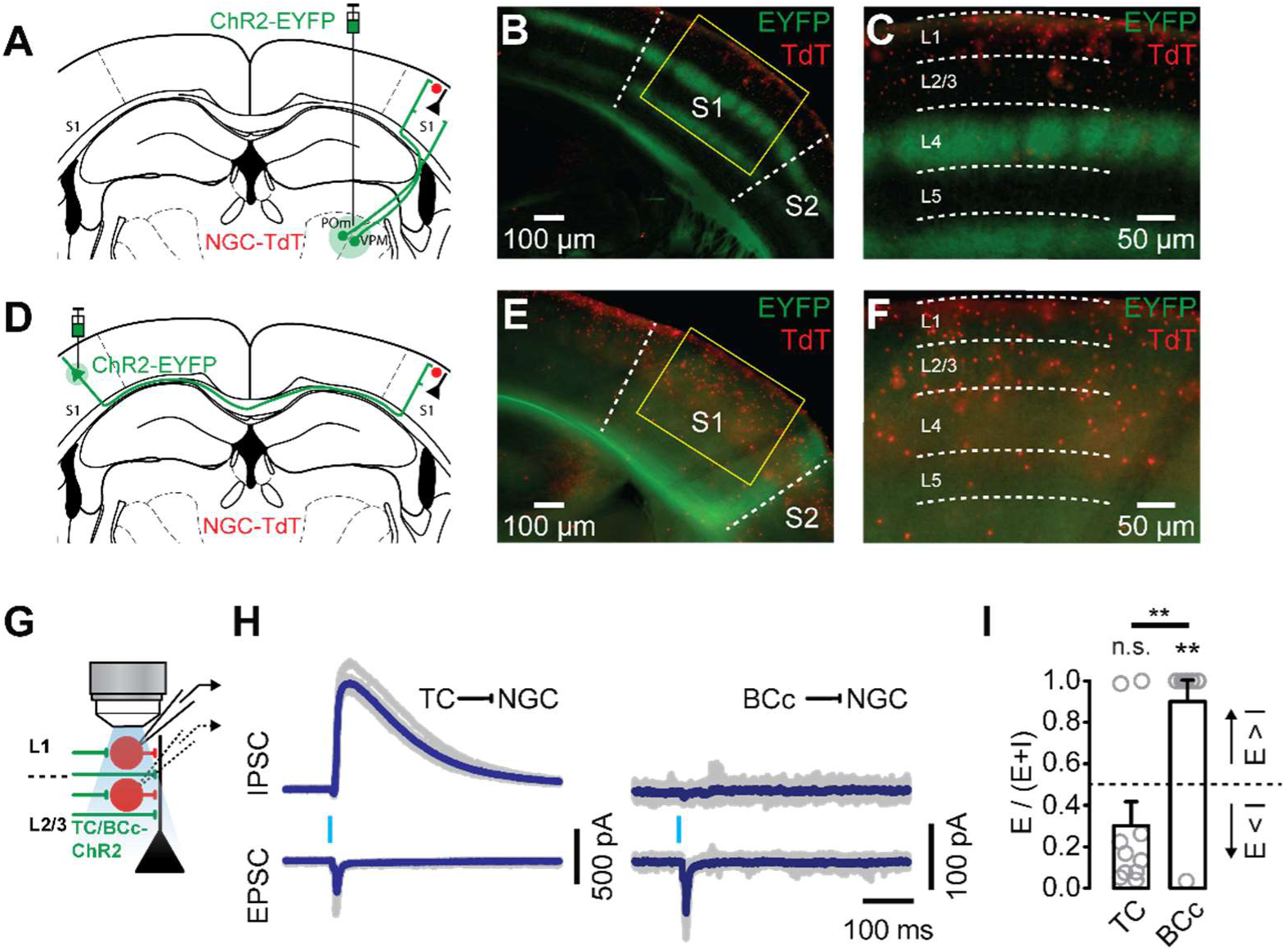
TC and BCc afferents drive inputs to L1-3 NGCs with differential E/I ratios. (A and D) Schematic representation of AAV-directed ChR2 expression in TC and BCc afferents, in NGC-TdT mice. (B and E) Low magnification epifluorescence images depicting the expression patterns in TC and BCc afferents in the BC of recorded slices. (C and F) Higher magnification epifluorescence images of boxed areas in B and E showing ChR2 expression in TC and BCc terminals, and TdT expression in L1-3 NGCs. (G) Schematic representation of whole-cell voltage clamp recordings from TdT-expressing L1-3 NGCs. (H) Representative examples of IPSCs (top) and EPSCs (bottom) recorded at Vh = 0 and Vh = −70mV respectively. Blue traces are the averages of individual traces shown in gray. (I) Comparison of E-I ratios between TC (*n=10* cells from *N = 3* mice) and BCc (*n=10* cells from *N = 3* mice) inputs. For TC vs BCc, **P = 4.7 x 10^-3^, Mann-Whitney test; forTC vs 0.5 and BCc vs 0.5, ^n.s.^P = 0.4316 and **P = 3.9 x 10^-3^, Wilcoxon rank sum test.

In voltage clamp recordings at −70 mV and 0 mV, optical stimulation of TC afferents evoked robust excitatory followed by inhibitory postsynaptic currents (EPSCs and IPSCs) in 80% of the NGCs, and only EPSCs in the other 20% (Figure 2H). In contrast, optical stimulation of BCc afferents evoked robust EPSCs followed by IPSCs in 10% of the cells, and only EPSCs in the other 90% (Figure 2H). Since, TC and BCc afferents are thought to be predominantly excitatory, we reasoned that the IPSCs were the result of polysynaptic feedforward connections. Indeed, when we bath-applied the glutamatergic blockers NBQX and AP5 to some of our voltage clamp recordings, we found that IPSCs were totally abolished, excluding the possibility that they were the result of monosynaptic input (Figure S2A-B). Conversely, when we performed current clamp recordings at resting membrane potentials, but in the presence of TTX and 4-AP (Figure S2C-D), optical stimulation of POm-derived TC as well as of BCc afferents evoked robust EPSPs in all cells (Figure S2C-D), providing proof that both projections constitute monosynaptic connections with NGCs. To quantify the differences in excitation and inhibition received from TC and BCc afferents, we computed the E-I ratio for each recorded cell by dividing its excitatory charge by the sum of its excitatory and inhibitory charges (Methods). Our analysis revealed a significantly higher E-I ratio of BCc compared to TC inputs (Figure 2I). Moreover, the E-I ratio of BCc inputs was significantly higher than 0.5 (Figure 2I). This could reflect the fact that excitation was higher than inhibition for BCc inputs, whereas it was lower than inhibition for TC inputs. We found very similar results when we analyzed L1 and L2/3 NGCs separately, meaning that the E-I ratio difference between TC and BCc inputs was not layer-specific (Figure S2F).

Altogether, these data indicate that both TC and BCc afferents provide monosynaptic excitation to L1-3 NGCs, but TC afferents provide also a much stronger inhibition in a feed-forward fashion, which is in line with our finding that responses of L2/3 PNs upon stimulation of TC afferents does not contain long-lasting inhibitory components (Figure 1D).

### L1-3 NGC activity is predominantly enhanced by ipsi- and suppressed by contra-lateral sensory inputs

We sought to validate the input and output connectivity of L1-3 NGCs *in vivo* in awake, head-restrained mice. First, we monitored the activity of L1-3 NGCs expressing the calcium sensor GCaMP6s in the BC, upon contra-, ipsi- or bi-lateral whisker stimulation (C-WS, I-WS, B-WS; Figure 3A), which primarily recruits TC, BCc, or both pathways, respectively. GCaMP6s was expressed in L1-3 NGCs using targeted injections of a flippase-dependent AAV in NGC-Flp mice (Methods). Successful expression in the center of the BC was validated by superimposing 2PLSM images on the barrel columns that had been mapped using intrinsic optical imaging (IOI; Methods). Single-cell calcium transients were monitored using two-photon laser scanning microscopy (2PLSM; Figure 3A). To simultaneously monitor the activity of as many NGCs as possible in L1 and L2/3, we performed fast volumetric imaging in two planes of extended depth of field ^24^ (Methods). Cells whose activity was significantly changed upon at least one of the whisker stimulation types across 60 trials, were defined as responding (93% of the total population; Methods). The rest were defined as non-responding and were excluded from our analysis (7% of the total population; Methods).

**Figure 3.**
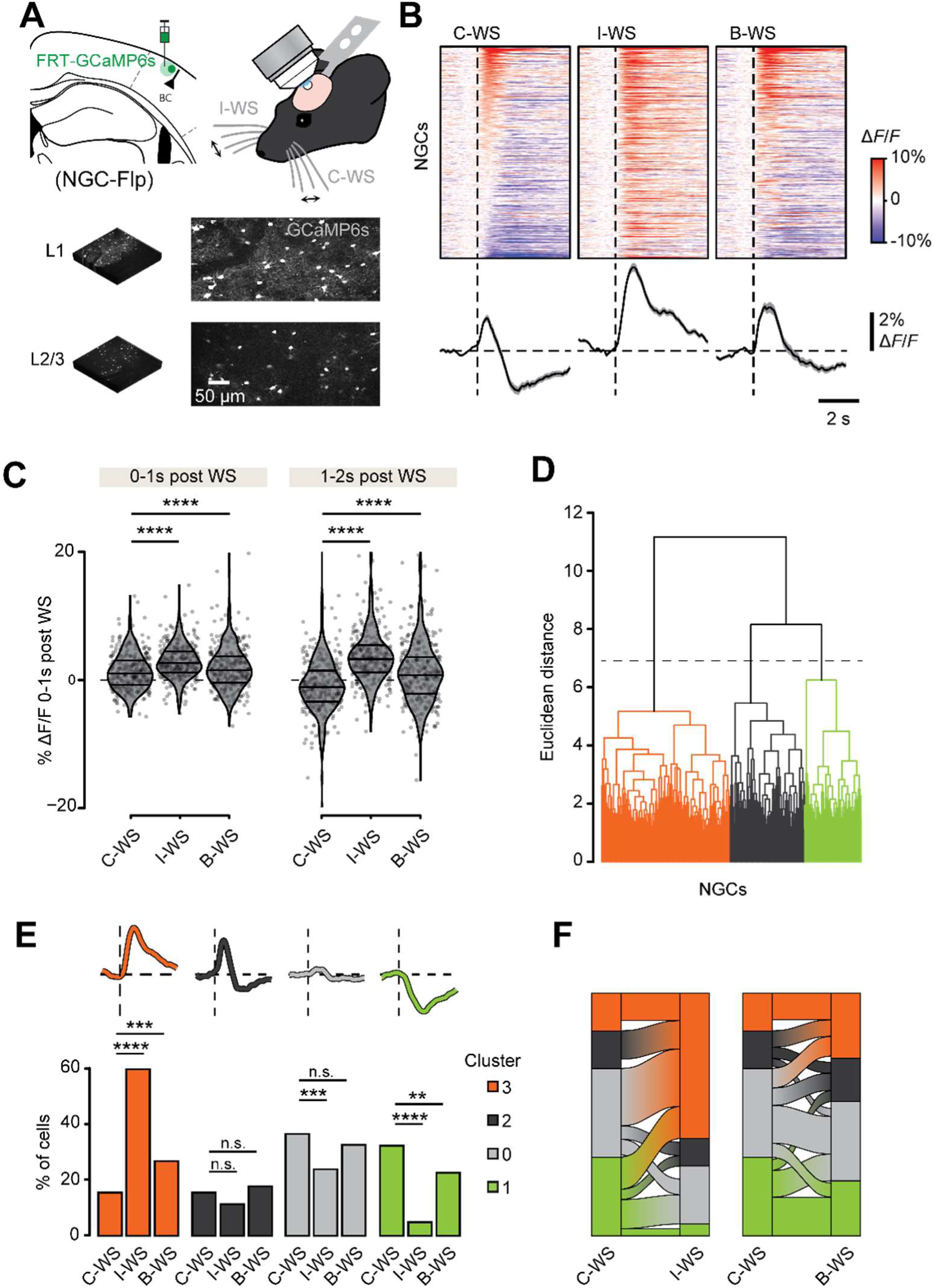
NGC activity is predominantly suppressed by contra- and enhanced by ipsi-lateral sensory inputs. (A) Schematic representation of calcium imaging in L1 and L2/3 NGCs upon WS. (B) Top, average calcium responses of each cell upon C-WS, I-WS, and B-WS, cross 60 trials. Responses are baseline corrected (subtraction of baseline Δ*F/F* −1 to 0 s before stimulus onset) and aligned to WS onset (dashed line). Cells are sorted according to their average response 0-1s post C-WS onset. Each plot includes *n=309* cells from *N=5* mice. Bottom, average response traces of all cells. Shaded areas represent S.E.M. (C) Comparison of average responses 0-1 s and 1-2 s post WS onset. C-WS vs I-WS for 0-1 s and 1-2 s post WS, ****P < 2.2 x 10^-16^ and ****P = 7.2 x 10^-7^; C-WS vs B-WS for 0-1 s and 1-2 s post WS, ****P < 2.2 x 10^-16^ and ****P < 2.2 x 10^-16^, paired t-test. (D) Hierarchical clustering dendrogram for the shape of responding cells’ average traces across all WS types. (E) Comparison of cluster proportions between different WS types. C-WS vs I-WS for clusters 3,2,0, and 1: ****P = 6.4 x 10^-31^, ^n.s.^P = 0.16, ***P = 8.5 x 10^-4^, and ****P = 1.1 x 10^-19^; C-WS vs B-WS for clusters 3,2,0, and 1: ***P = 7.8 x 10^-4^, ^n.s.^P = 0.52, ^n.s.^P = 0.35, and **P = 8.9 x 10^-3^; Fisher’s exact test of proportions. (F) Alluvial plots illustrating the proportions of cluster changes between stimulation types. Ribbons representing cluster changes for less than 2% of cells are removed for clarity.

The average activity of L1-3 NGCs was predominantly suppressed upon C-WS, enhanced upon I-WS, and bidirectionally modulated upon B-WS (Figure 3B). However, activity changes of individual NGCs were very heterogeneous for each stimulation type, with variable directions and time courses. To gain a deeper insight into different profiles of activity changes, we performed hierarchical clustering of the responding cells’ average traces (Figure 3D). We identified three major, functionally distinct profiles: one that showed a robust and sustained increase of activity in the 5 s post-stimulation window (cluster 3), one that showed an increase followed by a decrease (cluster 2), and one that showed a robust and sustained decrease (cluster 1) (Figure 3D and 3E). Traces of responding cells showing no change upon a specific stimulation type, were grouped separately, which we termed cluster 0 (Figure 3E). For I-WS compared to C-WS, the proportion of cells was significantly higher for cluster 3 and significantly lower for cluster 1 and cluster 0. Similar but less pronounced differences were observed also for B-WS compared to C-WS: the proportion of cells was significantly higher for cluster 3 and significantly lower for cluster 1. Interestingly, a large proportion of the cells in cluster 0, i.e. with a non-responding profile, upon C-WS, were part of cluster 3, i.e. with a sustained increase, upon I-WS (Figure 3F). This suggests that many of the NGCs that respond to I-WS are not part of a C-WS synaptic pathway, or are part of a C-WS synaptic pathway in which excitation and inhibition cancel each other out. This is in line with the electrophysiology data showing that only 20% of the NGCs depolarize upon TC stimulation, whereas 80% does so upon BCc stimulation.

To test whether these results varied between L1 and L2/3 NGCs, we also analyzed these two groups separately. Interestingly, we found that the suppression upon C-WS was more pronounced in L1 NGCs, whereas the enhancement upon I-WS was more pronounced in L2/3 NGCs (Figure S3A-B). Interestingly, we also found that activation and suppression upon B-WS were more pronounced in L2/3 and L1 respectively. Similarly, the proportion of cells in cluster 1 upon C-WS was significantly higher for L1 NGCs, whereas the proportion of cells in clusters 3 upon I-WS was higher, though not significantly, for L2/3 NGCs (Figure S3C).

Overall, these data indicate that NGC activity is predominantly enhanced by I-WS and suppressed by C-WS, and that these opposing effects are more pronounced in L2/3 and L1 NGCs respectively, implying that apical dendrites of PNs running through L2/3 could receive more pronounced inhibition from NGCs upon sensory stimuli as compared to their tufts in L1.

### Optogenetic activation of NGCs suppresses sensory-evoked activity in L2/3 PNs, but this is only partially recapitulated by ipsilateral sensory inputs

Next, we probed the net modulatory effect of L1-3 NGCs on sensory-evoked activity in PNs. First, we monitored the activity of L2/3 PNs expressing the calcium sensor GCaMP6s in the BC (Figure 4A) and tested how C-WS-evoked responses were modulated upon pairing the C-WS with optical stimulation of ChR2-expressing L1-3 NGCs (C-WS&LED) or with I-WS (B-WS). ChR2 was expressed in L1-3 NGCs using targeted injections of a flippase-dependent AAV in NGC-Flp mice (Methods). GCaMP6s was expressed in L2/3 PNs using targeted injections of a constitutive AAV carrying the CaMKII promoter. Successful expression of ChR2 and GCaMP6s in the center of the BC was validated by superimposing 2PLSM images and the barrel map that was defined with intrinsic optical imaging (IOI; Methods). Single-cell calcium transients were monitored using 2PLSM in one plane at the level of L2/3 (Figure 4A). Cells whose activity was significantly enhanced upon C-WS across 60 trials, were defined as responding (61% of the total population). The rest were defined as non-responding and were excluded from our analysis (Methods).

**Figure 4.**
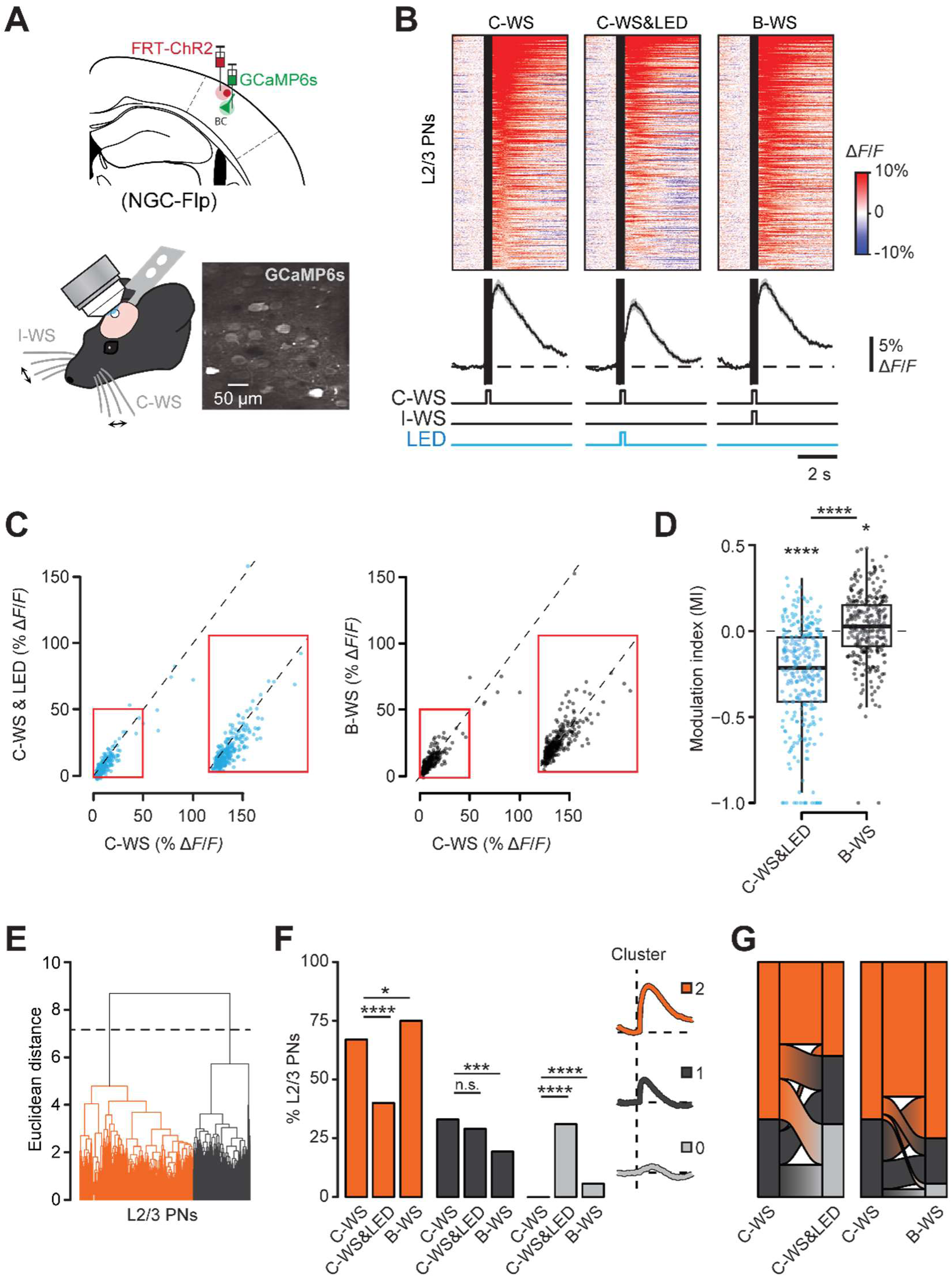
NGC activation and I-WS modulate sensory-evoked activity in L2/3 PNs. (A) Schematic representation of calcium imaging in L2/3 PNs upon WS combined with optogenetic activation of L1-3 NGCs. (B) Top, average calcium responses of each cell upon C-WS, C-WS&LED, and B-WS, across 60 trials. Responses are baseline corrected (subtraction of baseline Δ*F/F* −1 to 0 s before stimulus onset) and aligned to WS onset (black band left border). Black bands indicate periods of no data collection due to the photomultiplier tubes’ shutdown during LED stimulation. Cells are sorted according to their average response 0-2s post C-WS onset. Each plot includes *n = 300* cells from *N = 3* mice. Bottom, average response traces of all cells. Shaded areas represent S.E.M. (C) Scatter plots of average responses 0-2s post WS for C-WS vs C-WS&LED and C-WS vs B-WS. (D) Comparison of C-WS&LED and B-WS modulation indexes with zero and between them. C-WS&LED vs zero and B-WS vs zero: ****P = 8.3 x 10^-35^ and *P = 0.011, Wilcoxon signed rank test; C-WS&LED vs B-WS: ****P = 5.4 x 10^-30^; Wilcoxon rank sum test. (E) Hierarchical clustering dendrogram for the shape of responding cells’ average traces across all WS types. (F) Comparison of cluster proportions between different WS types. C-WS vs C-WS&LED for clusters 2,1, and 0: ****P = 4.5 x 10^-11^, ^n.s.^P = 0.33, ****P = 4.2 x 10^-32^; C-WS vs B-WS for clusters 2,1, and 0: *P = 0.038, ***P = 1.9 x 10^-4^, ****P = 1.2 x 10^-5^, Fisher’s exact test. (G) Alluvial plots illustrating the proportions of cluster changes between stimulation types.

In contrast to the asynchronous bilateral movement of paws, bilateral whisking is mostly synchronous^25^. Hence, we used a protocol of synchronous pairing. On average, C-WS-evoked activity was reduced upon synchronous pairing of C-WS with optical stimulation of NGCs (C-WS&LED), but it was slightly increased upon synchronous pairing of C-WS with I-WS (B-WS; Figure 4B-C). To quantify the modulatory effect of C-WS&LED and B-WS, we computed a modulation index (MI) for each neuron ^26^, in both pairing regimes. C-WS&LED MIs were much lower and significantly different from zero, whereas B-WS MIs were slightly higher but also significantly different from zero (Figure 4D). Moreover, C-WS&LED and B-WS MIs were significantly different between them (Figure 4D).

Upon the hierarchical clustering of the responding cells’ average traces (Figure 4E), we identified two major, functionally distinct profiles: One showing a robust and prolonged increase of activity in the 5 s post-stimulation window (cluster 2), and one showing a moderate and short increase (cluster 1). Traces of responding cells showing no change upon a specific stimulation type, were grouped as cluster 0 (Figure 4F). For C-WS&LED compared to C-WS, the proportion of cells was significantly lower for cluster 2 and significantly higher for cluster 0 (Figure 4F). The strong reduction in the fraction of cluster 2 cells upon C-WS&LED was due to a transfer of approximately a quarter of the cells to cluster 1 and another quarter to cluster 0 (Figure 4G). However, we observed the opposite differences for B-WS compared to C-WS: the proportion of cells was significantly higher for cluster 0, due to a relatively large transfer of cells from cluster 1 in C-WS to cluster 2 in B-WS, accompanied by a relatively small transfer of cells from cluster 2 in C-WS to other clusters in B-WS (Figure 4G). The proportion of cells in cluster 0 upon B-WS was also significantly higher than in C-WS, but it was very small (Figure 4F), suggesting that I-WS did not dampen C-WS-evoked activity. Indeed, the activity that we observed upon B-WS was supported by the observation that I-WS alone did not suppress but rather enhance the overall L2/3 PN activity, although to a lesser extent as compared to C-WS (Figure S4A-D). Indeed, when comparing C-WS to I-WS, we found only about a quarter of the cells in cluster 2 cells in C-WS to transfer to cluster 0 in I-WS (Figure S4E).

Overall, these data indicate that C-WS-evoked activity in L2/3 PNs is reduced upon synchronous pairing of C-WS with optical stimulation of NGCs. However, this effect is not recapitulated upon synchronous pairing of C-WS with I-WS, most likely due to the fact that I-WS alone enhances activity of the same cells that display enhanced activity upon C-WS. The data also indicate that the cells whose activity is not enhanced by I-WS alone, are neither strongly inhibited by it. This suggests that the NGCs that are activated by I-WS are not efficiently suppressing B-WS-evoked activity of L2/3 PNs.

### L5 PN activity is suppressed when NGC stimulation or ipsi-lateral sensory inputs precede the contralateral sensory stimulus

We asked the same questions for L5b PNs. For this, we monitored the activity of L5b PN apical tufts expressing the calcium sensor GCaMP7s in the BC, and performed the same manipulations and analysis as described for L2/3 PNs (Figure 5A). GCaMP7s was expressed in L5b PNs retrogradely using targeted injections of a constitutive rgAAV in the superior colliculus (SC; Figure 5A). Calcium transients in apical dendritic tufts were monitored using 2PLSM in one plane at the level of L1 (Figure 5A).

**Figure 5.**
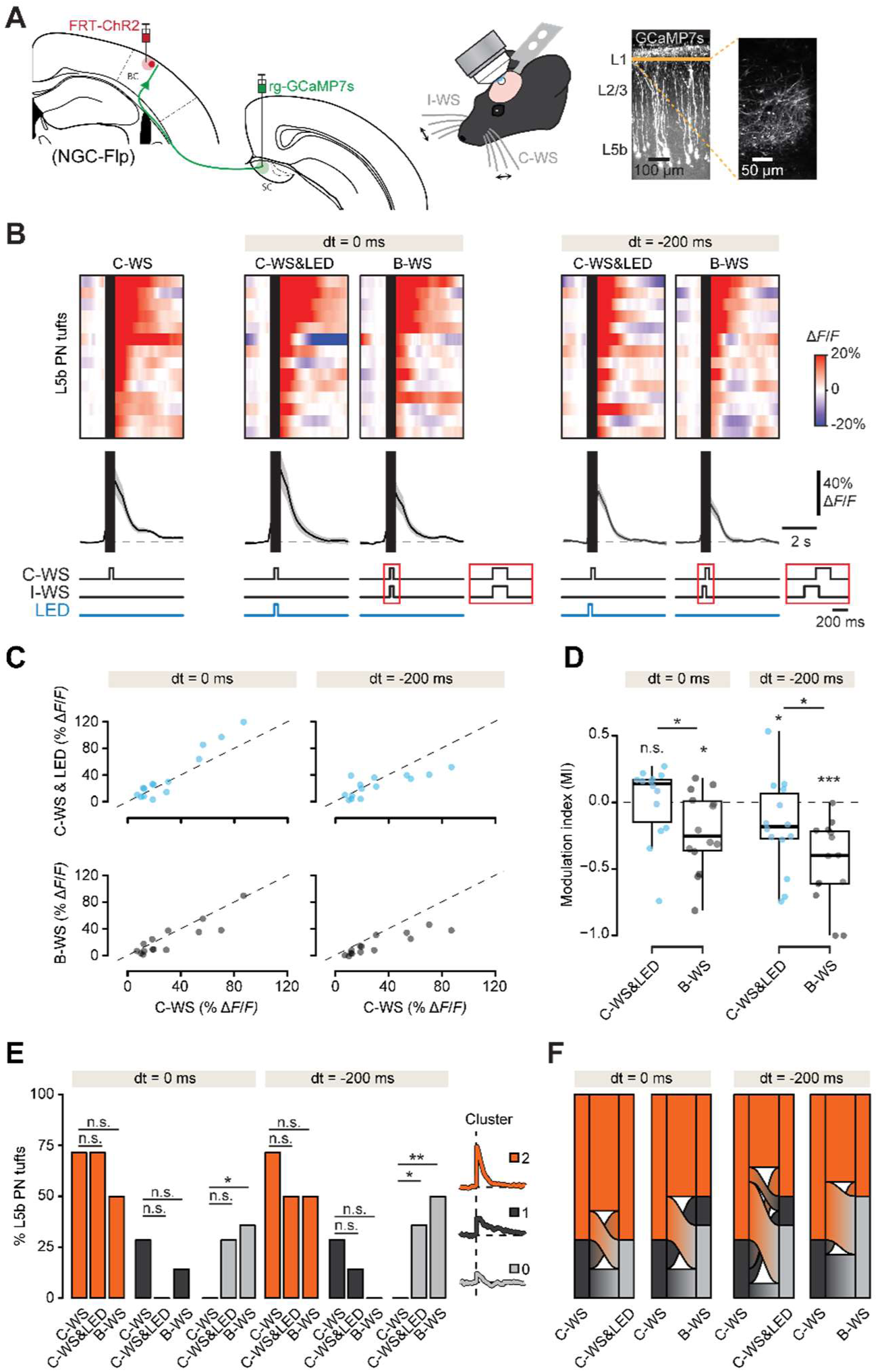
NGC activation and I-WS modulate sensory-evoked activity in L5b PN tufts. (A) Schematic representation of calcium imaging in L5b PN tufts upon WS combined with optogenetic activation of L1-3 NGCs. (B) Top, average calcium responses of each tuft to C-WS, C-WS&LED, and B-WS, across 60 trials, with 0 or −200 ms delay in pairing. Responses are baseline corrected (subtraction of baseline Δ*F/F* −1 to 0 s before stimulus onset) and aligned to WS onset (dark band left border). Tufts are sorted according to their average response 0-1s post C-WS onset. Each plot includes *n = 14* tufts from *N = 2* mice. Bottom, average response traces of all tufts. (C) Scatter plots of average responses 0-2s post WS for C-WS vs C-WS &LED and C-WS vs B-WS, with 0 or −200 ms delay in pairing. (D) Comparison of C-WS&LED and B-WS modulation indexes with zero and between them, for 0 or −200 ms delay in pairing. C-WS&LED vs zero and B-WS vs zero: ^n.s.^P = 0.71 and *P = 0.024 for dt = 0 ms; *P = 0.042 and ***P = 1.2 x 10^-4^ for dt = −200 ms, Wilcoxon signed rank test; C-WS&LED vs B-WS: *P =0.049 for dt = 0 ms; *P =0.035 for dt = −200 ms, Wilcoxon rank sum test. (E) Comparison of cluster proportions between different WS types. C-WS vs C-WS&LED for clusters 2,1, and 0: ^n.s.^P = 1, ^n.s.^P = 0.098, ^n.s.^P = 0.098 for dt = 0 ms; ^n.s.^P = 0.44, ^n.s.^P = 0.65, *P = 0.041 for dt = −200 ms; C-WS vs B-WS for clusters 2,1, and 0: ^n.s.^P = 0.44, ^n.s.^P = 0.65, *P = 0.041 for dt = 0 ms; ^n.s.^P = 0.44, ^n.s.^P = 0.098, **P = 5.8 x 10^-3^ for dt = −200 ms; Fisher’s exact test. (G) Alluvial plots illustrating the proportions of cluster changes between stimulation types.

Surprisingly, we observed the opposite results as compared to L2/3 PNs upon synchronous pairing regimes: on average, C-WS-evoked activity was slightly increased upon synchronous pairing of C-WS with optical stimulation of NGCs (C-WS&LED, dt = 0 ms), and decreased upon synchronous pairing of C-WS with I-WS (B-WS, dt = 0 ms; Figure 5B-C). C-WS&LED MIs were slightly higher but not significantly different from zero, whereas B-WS MIs were much lower and significantly different from zero (Figure 5D, dt = 0 ms). Moreover, C-WS&LED and B-WS MIs were significantly different between them (Figure 5D, dt = 0 ms).

To test whether the modulation of C-WS-evoked responses by NGC optical stimulation and I-WS depends on the timing of pairing, we employed a protocol of asynchronous pairing, in which optogenetic stimulation of NGCs or I-WS preceded the C-WS by 200 ms (dt = −200 ms). A similar protocol was described in previous work on interhemispheric inhibition in the hindlimb area of S1 (S1HL) ^7^. On average, C-WS-evoked activity was decreased upon both asynchronous pairing regimes (C-WS&LED and B-WS; dt = −200 ms; Figure 5B-C). Both C-WS&LED MIs and B-WS MIs were much lower and significantly different from zero, (Figure 5D, dt = −200 ms), and significantly different between them (Figure 5D, dt = −200 ms). In both cases, a large portion of the cluster 2 cells in C-WS (close to half) transferred to cluster 0 in C-WS&LED or B-WS (Figure 5F), which is supported by finding that I-WS alone evokes a negligible increase of the calcium signals in L5b PN tufts (Figure S5A-E). Altogether, these data indicate that that C-WS-evoked activity in L5b PN apical tufts is suppressed when optical stimulation of NGCs precedes the C-WS. This effect is recapitulated when I-WS precedes C-WS. Notably, the modulatory effect of I-WS is even stronger as compared to NGC stimulation. Together, this suggests that the activation of NGCs through I-WS is able to suppress B-WS-evoked responses in L5b PN apical tufts. In turn, this suggests that I-WS may interfere with cortical processing of the C-WS, through NGC-mediated modulation of L5b PNs, which convey cortical outputs to subcortical areas.

### Optogenetic stimulation of L1-3 NGCs and ipsi-lateral sensory inputs increase the perceptual threshold in a whisker-stimulation detection task

Based on our observations above, we hypothesized that NGC-mediated modulation of sensory-evoked activity in PNs affects sensory perception. To answer this, we employed a single whisker stimulation detection task ^27,28^ and tested how pairing the C2 whisker stimulation (C2S) with optical stimulation of NGCs (C2S&LED) or I-WS (C2S&I-WS) modulates sensory perception. Mice were trained to report whisker stimulations by licking a liquid dispenser to collect sugar water rewards (Figure 6A). Selective single whisker stimulation was achieved by inserting the left snout’s C2 whisker in a hollow metallic bar and delivering magnetic pulses through a magnetic coil placed under the mouse’s head (Methods). For each mouse, after learning the task upon salient stimulation, we computed its perceptual detection threshold from a psychometric function based on the detection probabilities at variable stimulation intensities ^27–29^ (Figure 6B,E; Methods). ChR2 was expressed in L1-3 NGCs using targeted injections of a flippase-dependent AAV in NGC-Flp mice (Methods). Successful expression of ChR2 in the center of the BC was validated by superimposing 2PLSM images onto the barrel map that was determined using intrinsic optical imaging (IOI; Methods). To verify that whisker stimulus perception at variable intensities critically depends on activity in S1, we used a mouse line in which ChR2 expression is directed to all interneurons under control of the vesicular inhibitory amino acid transporter (VGAT) promoter (VGAT-ChR2; Figure S6). VGAT-ChR2 mice are commonly used to efficiently suppress S1 BC activity ^30–32^. We used a protocol of asynchronous pairing in which the optogenetic stimulation of NGCs preceded the C-WS by 200 ms (dt = −200 ms), similar to the protocol that efficiently modulated sensory-evoked activity in L5b PN tufts (Figure 5).

**Figure 6.**
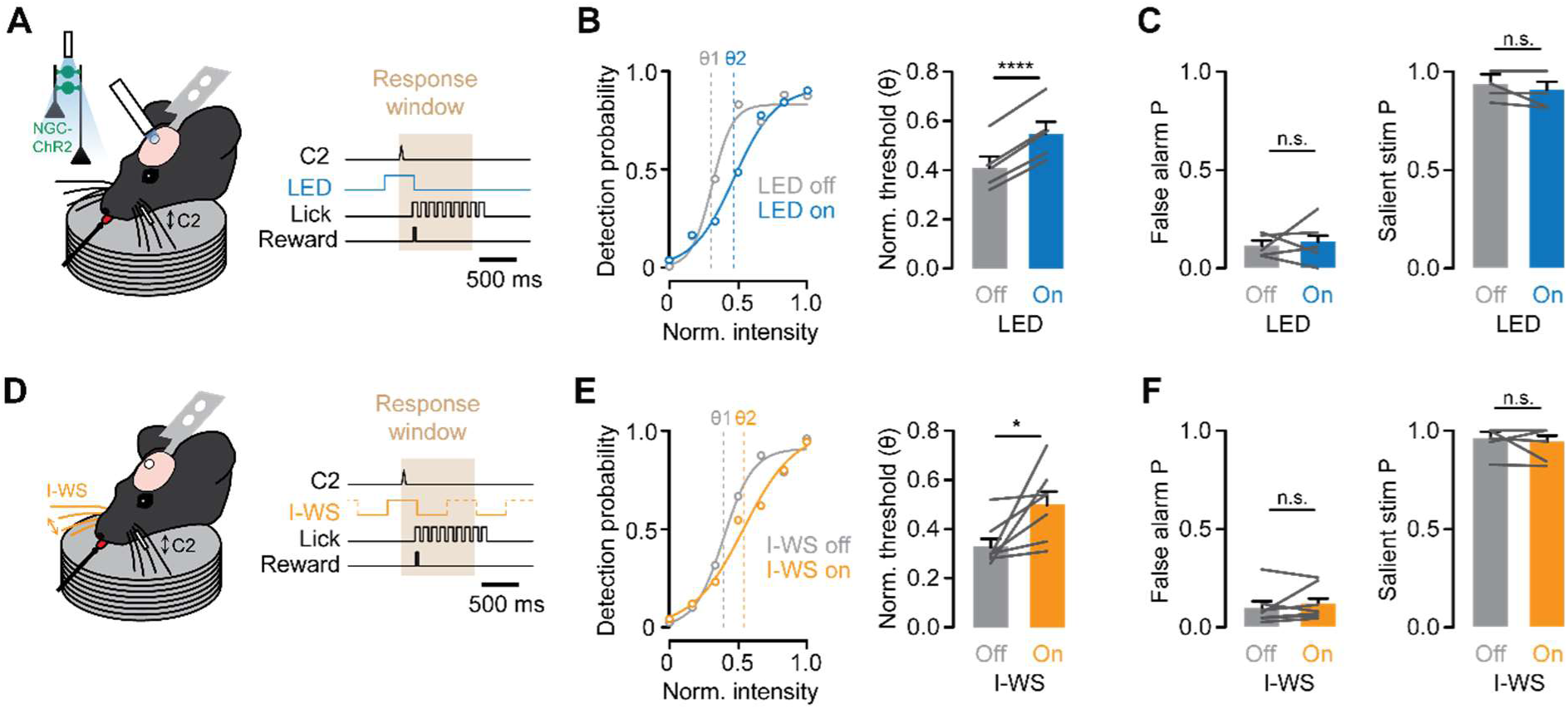
Direct optical and indirect sensory activation of L1-3 NGCs increase the perceptual threshold in a whisker-deflection detection task. (A and D) Schematic representation of the behavioral task combined with optical stimulation of ChR2-expressing L1-3 NGCs or I-WS. (B and E) Left, representative examples of psychometric curves for trials without manipulation (gray) and trials with optical stimulation of L1-3 NGCs (blue) or I-WS (orange). Right, normalized detection thresholds (Θ) for trials without manipulation (gray) and trials with optical stimulation of L1-3 NGCs (blue; *N = 5* mice) or I-WS (orange; *N = 7* mice). ****P = 6.5 x 10^-5^ for B, and *P = 0.0476 for E, paired t-test. (C and F) Detection probabilities for trials without manipulation (gray) and trials with optical stimulation of L1-3 NGCs (blue; *N = 5* mice) or I-WS (orange; *N = 7* mice) at zero stimulation intensity (false alarm) and salient stimulation intensity. ^n.s.^P = 0.7420 and ^n.s.^P = 0.2795 for C; ^n.s.^P = 0.4010 and ^n.s.^P = 0.5124 for F, paired t-test.

The detection threshold was significantly increased upon pairing of C2S with optical stimulation of the general population of interneurons (Figure S6; C2S&LED vs C2) as well as by NGCs in particular (Figure 6B; C2S&LED vs C2). Consistent with our hypothesis, this was recapitulated upon pairing of C2S with I-WS (C2S&I-WS; Figure 6E). Notably, we found no significant differences between C2S and C2S&LED, nor between C2S and C2S&I-WS for the 0 and 1.0 stimulus intensities (Figure 6C, F). This rules out that the perceptual effects were biased due to a possible distraction of the mice caused by the LED or the I-WS, since this would have also led to an increase in false alarms and/or a decrease in the salient stimulus detection probabilities. Together, these data demonstrate that direct activation of NGCs modulates sensory perception, and that this effect is recapitulated by I-WS. This suggests that the transcallosal pathway may interfere with sensory perception through the activation of NGCs, which suppresses the sensory-evoked activity of L5b PNs.

## DISCUSSION

NGCs have long been hypothesized to mediate interhemispheric inhibition of cortical PNs ^7^. However, direct evidence supporting this hypothesis was missing due to a lack of genetic access to NGCs. Here, we used selective genetic targeting, *ex vivo* whole cell recordings, optogenetics, *in vivo* 2-photon calcium imaging, and perceptual detection behavior, to unravel the input and output connectivity of NGCs, and to provide direct evidence for their activation by interhemispheric callosal afferents to modulate sensory processing and perception.

### NGC input and output connectivity

We showed that optical stimulation of BCc but not TC afferents drives a GABA_B_R-dependent inhibition of L2/3 PNs (Figure 1), which corroborates and expands previous findings of transcallosal GABA_B_R-dependent inhibition of L5 PNs ^7^. The results align with other studies showing that optical stimulation of NDNF INs, which comprise L1 NGCs ^10,33^, elicits GABA_A_R and GABA_B_R-dependent inhibition in the neocortex. By leveraging the *Hmx3*-Cre mouse line, we were able to reveal and quantify the specific contribution of NGCs to the GABA_A_R- and GABA_B_R-dependent components of the inhibition in L2/3 and L5b PN somata, as well as in L5b PN dendrites. Using this mouse line we also demonstrated that L1-3 NGCs receive both monosynaptic excitation and polysynaptic inhibition from TC afferents, yet only monosynaptic excitation from BCc afferents (Figure 2). This is in agreement with previous studies in L1 NDNF INs, ∼30% of which are NGCs ^16^, showing similar monosynaptic inputs from TC afferents in the A1 and V1 ^11,13,14^, and from BCc afferents in the S1 ^12^. Hence, our data proves that the current knowledge on monosynaptic excitation of L1 NDNF INs holds true also for L1 NGCs, and expands this knowledge to L2-3 NGCs as well. Moreover, recent studies have shown that L1 NDNF INs are under the inhibitory control of SST but not of PV or VIP INs ^10,33^. Therefore, it is very likely that the TC afferent-induced polysynaptic inhibition shown in our experiments arises from recruitment of SST INs which in turn inhibit L1-3 NGCs in a feed-forward fashion. This feedforward polysynaptic inhibition of NGCs by TC afferent input may have the ability to cancel out, at least partially, the excitatory inputs to NGCs from this pathway, which aligns with the observation that responses of L2/3 PNs upon stimulation of TC afferents does not contain obvious long-lasting inhibitory components (Figure 1).

### Role of direct and indirect stimulation of NGCs in sensory processing

The activity of NGCs and PNs upon sensory-mediated engagement of the TC and BCc pathways in awake head-restricted mice confirmed our *ex vivo* results. We showed that NGC activity is largely suppressed by C-WS and predominantly enhanced by I-WS, and that these responses partially cancel each other out upon B-WS (Figure 3). Previous studies have shown that L1 NDNF INs in V1 are activated upon contralateral sensory stimulation ^11^, and that ∼40% of L1 INs in S1 are activated upon ipsilateral sensory stimulation ^7^. However, these studies included only cells showing positive responses in calcium imaging. Our data shows that sensory stimulation can change baseline activity of L1-3 NGCs bidirectionally, revealing mixed responses within and between cells. It demonstrates that the previously observed enhancement of activity in the L1 NDNF IN population upon contralateral sensory stimulation holds true only for a small fraction of L1 NGCs, and expands this further to a fraction of L2-3 NGCs as well. Interestingly, we also found that the suppression of NGC activity by C-WS is more pronounced in L1, in contrast to the enhancement by I-WS, which was more pronounced in L2/3 NGCs (Figure S3). This suggests that slow interhemispheric inhibition is mediated by both L1 and L2/3 NGCs, but most likely relies more on the L2/3 NGCs.

We assessed the impact of NGC-induced inhibition on sensory-evoked activity of PNs. C-WS-evoked activity in L2/3 PNs was reduced upon synchronous pairing of C-WS with optogenetic stimulation of L1-3 NGCs (Figure 4), which is consistent with observations in a study using optical stimulation of L1 NDNF INs in V1 ^11^. Asynchronous pairing of C-WS with optical stimulation of L1-3 NGCs also reduced C-WS-evoked activity in L5b PN distal dendrites (Figure 5), which is consistent with observations in a study using optical stimulation of L1 NDNF INs in A1 ^10^. What is more, we showed that C-WS-evoked activity in L5b PNs was reduced upon both synchronous and asynchronous pairing of C-WS with I-WS (Figure 5). This is also in agreement with previous studies in S1HL ^7^ and in BC ^12^. Since I-WS strongly activates NGCs through BCc afferents (Figure 3), this finding suggests that, similar to optical stimulation of NGCs, pairing C-WS with I-WS (i.e. B-WS) reduces sensory-evoked activity of PNs by counteracting on the suppression of L1-3 NGCs that normally would happen upon C-WS (Figure 3). Altogether, this constitutes strong evidence for the notion that I-WS interhemispheric inhibition of PNs is largely mediated by NGCs. The ultimate experiment to causally link the modulatory effect of I-WS with NGC activation, would be to combine I-WS with optical silencing of NGCs. However, this is challenging due to the incompatibility of calcium imaging with long optical pulses needed for fast reversible silencing of neurons with commonly used optogenetic probes.

### Role of direct and indirect stimulation of NGCs in perceptual detection

Last, we interrogated the impact of NGC-induced inhibition on sensory perception. We showed that asynchronous pairing of C2-WS with optical stimulation of L1-3 NGCs increased the perceptual threshold in a whisker-deflection detection task (Figure 6). This effect was recapitulated upon asynchronous pairing of C2-WS with I-WS. These findings are consistent with previous observations that sensory perception in the BC depends on L5b PN distal dendritic activity, and is sensitive to pharmacological activation of GABA_B_Rs ^27,28^. Based on our data on NGC output connectivity and the role of NGCs in sensory perception, our results provide strong evidence that activation of L1-3 NGCs upon optical stimulation or sensory-mediated engagement of BCc afferents, modulates sensory perception mainly via suppression of L5b PN distal dendrite activity, in a GABA_B_R-dependent fashion. To determine the degree to which the effect of interhemispheric inhibition on sensory perception depends on NGCs and/or GABA_B_Rs, future experiments should combine I-WS with silencing the activity of NGCs and/or pharmacological blockage of GABA_B_Rs. In the context of sensory perception, it is also interesting to note that that monosynaptic excitation of NGCs from TC afferents may predominantly arise from the POm (Figure S2). This suggests that the POm inputs may counteract on the polysynaptic inhibition of NGCs. The POm is a higher-order sector of the somatosensory thalamus that is implicated in contextual feedback, and most strongly activated during active whisking-mediated sensory perception rather than upon passive whisker stimuli ^34,35^. Thus, the activation of NGCs by the Pom afferents may critically contribute to switching the L1 population of NGCs from inactive to active during active-sensory perception by overcoming the dominating polysynaptic inhibition of NGCs and thereby facilitating the transcallosal inhibition of PNs.

## STAR METHODS

### Key resources table

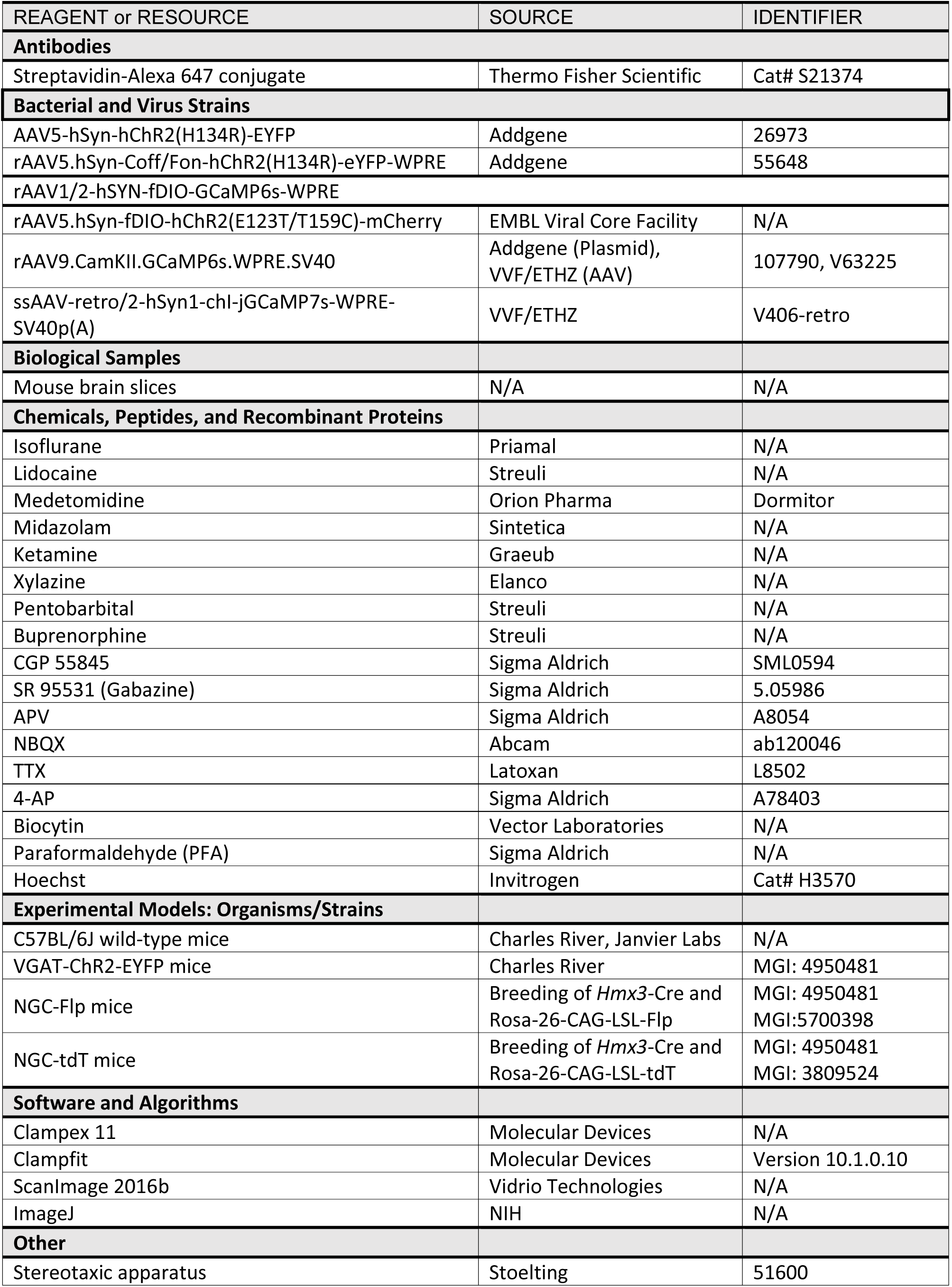
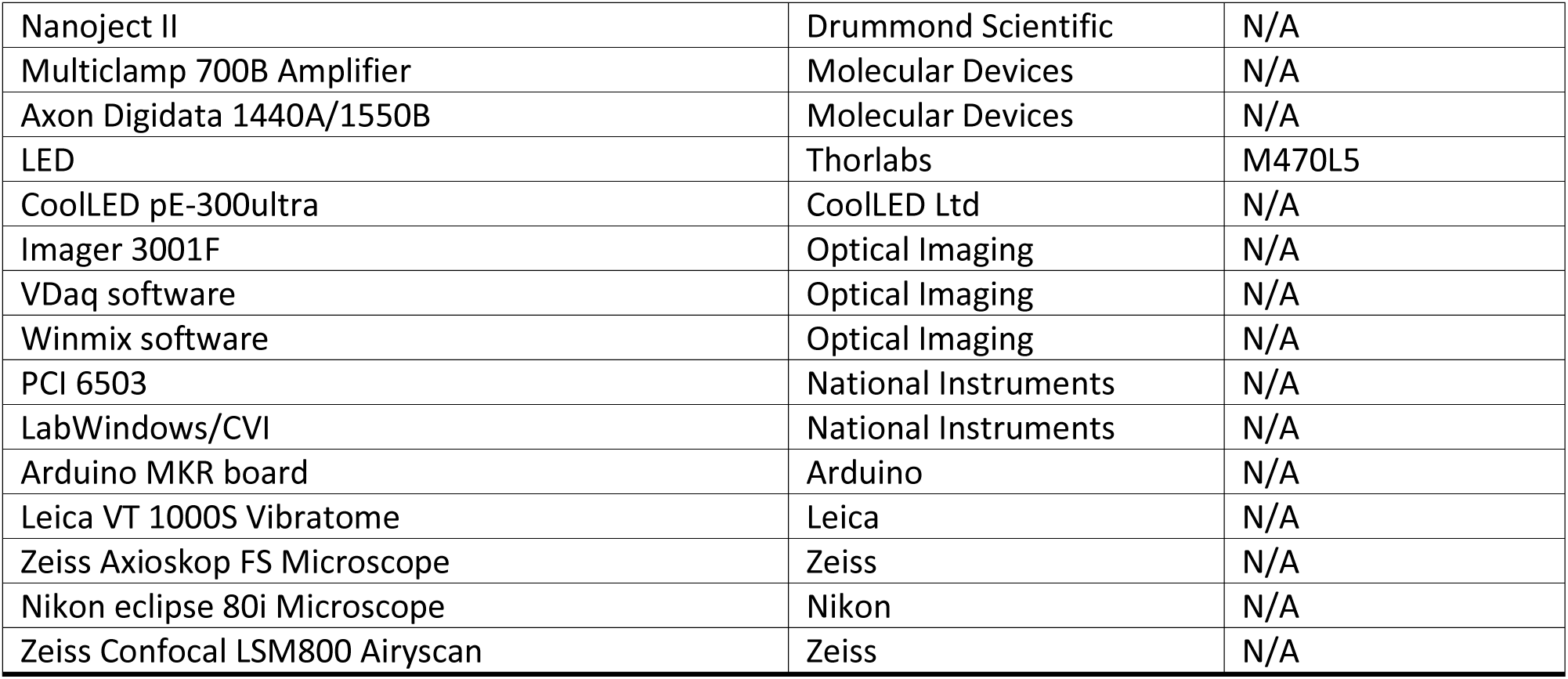

## CONTACT FOR REAGENT AND RESOURCE SHARING

Information and request for reagents may be directed and will be fulfilled by the Lead Contact Anthony Holtmaat.

## EXPERIMENTAL MODEL AND SUBJECT DETAILS

### Animals

Male and female C57BL/6J wild-type mice (Charles River, Janvier Labs or born in house), VGAT-ChR2-EYFP mice (MGI: 4950481 https://www.jax.org/strain/014548, RRID: IMSR_JAX:014548), NGC-Flp mice – obtained from the breeding of *Hmx3*-Cre (https://www.alliancegenome.org/allele/MGI:5566775) and Rosa-26-CAG-LSL-Flp (http://www.informatics.jax.org/allele/MGI:5700398), and NGC-tdT mice – obtained from the breeding of *Hmx3*-Cre and Rosa-26-CAG-LSL-tdT (MGI: 3809524 http://www.informatics.jax.org/allele/key/63704), aged 8 to 12 weeks, were group housed with littermates on a normal 12-h light cycle with food and water available *ad libitum*. All procedures were conducted in accordance with the guidelines of the Federal Food Safety and Veterinary Office of Switzerland and in agreement with the veterinary office of the Canton of Geneva (license numbers GE12219B, GE/74/18 and GE253A).

## METHODS DETAILS

### Virus injections and implantation of cranial windows

Mice of 8–12 weeks old, were anesthetized with isoflurane mixed with O_2_ (3–5% for induction, 1–2% for maintenance), placed in a stereotaxic apparatus, and prepared for injections with craniotomies over the target injection regions. Deep anesthesia was assessed by absence of foot pinch reaction. The skin overlying the skull was incised or removed under local anesthesia using Lidocaine (Streuli). Mice were then head-fixed with ear-bars and a nose clamp on a stereotaxic apparatus (Stoelting). Eyes were protected from drying with artificial tears. The body temperature was monitored with a rectal probe and was maintained at ∼37°C using a heating pad (FHC) during surgery. Unilateral or bilateral craniotomies were performed using an air-pressurized driller and injections (100-200 nl per injection site) were performed using a pulled glass pipette (10–15 µm diameter tip) mounted on a Nanoject II small-volume injector (Drummond Scientific). Injections were performed at a speed of 23 nl/s, separated by 2-3 min intervals.

For *ex vivo* electrophysiology: To express ChR2 in TC or BCc afferents, AAV5-hSyn-hChR2(H134R)-EYFP (Addgene, 26973) was injected in POm (2.2 mm posterior to bregma, 1.2 mm lateral and 3 mm below the bregma) and VPM (1.85 mm posterior to bregma, 1.75 mm lateral and 3.5 mm below the bregma) or S1 (1.5 mm posterior to bregma, −3.5 mm lateral and 0.4 mm below the pial surface), in NGC-tdT or WT mice. To express ChR2 in NGCs, rAAV5.hSyn-Coff/Fon-hChR2(H134R)-eYFP-WPRE (Addgene, 55648) was injected in S1 (1.5 mm posterior to bregma, 3.5 mm lateral and 0.2 mm below the pial surface), in NGC-Flp mice.

For calcium imaging and behavior: To express GCaMP or ChR2 in NGCs, rAAV1/2-hSYN-fDIO-GCaMP6s-WPRE (EMBL viral core facility, Rome, Italy), rAAV5.hSyn-fDIO-hChR2(E123T/T159C)-mCherry (EMBL viral core facility, Rome, Italy), or rAAV5.hSyn-Coff/Fon-hChR2(H134R)-eYFP-WPRE (Addgene, 55648) was injected in S1 (1.5 mm posterior to bregma, 3.5 mm lateral and 0.2 mm below the pial surface) in NGC-Flp mice. To express GCaMP in L2/3 or L5b PNs, rAAV9.CamKII.GCaMP6s.WPRE.SV40 (Plasmid from Addgene, 107790, AAV production from VVF/ETHZ, V63225) was injected in S1 (1.5 mm posterior to bregma, 3.5 mm lateral and 0.4 mm below the pial surface), or ssAAV-retro/2-hSyn1-chI-jGCaMP7s-WPRE-SV40p(A) (AAV production from VVF/ETHZ, V406-retro) was injected in SC (3.5 mm posterior to bregma, 1.25 mm lateral and 1.70 mm below the bregma), in NGC-Flp mice. A 3-mm diameter cranial window was implanted above S1, as described previously ^36^. Following this procedure, a metal post was implanted laterally to the window using dental acrylic to restrict head movement during imaging and behavior.

### Intrinsic optical imaging

Two weeks post surgery, three central barrel columns (typically the C2, C1, D1) were mapped using intrinsic optical imaging to validate the location of GCaMP and/or ChR2 expression in the barrel cortex, for Ca^2+^ imaging and/or behavioral experiments. For this, mice were anesthetized using induction with isoflurane (4%; 0.4 L min^−1^) followed by an intraperitoneal injection of a medetomidine/midazolam (0.2 mg kg^−1^/5 mg kg^−1^) mix in 0.9% NaCl. Each individual whisker was inserted into a capillary connected to a piezo actuator. Intrinsic signals were collected upon repeated whisker stimulations (1 s at 8 Hz). A 100-W halogen light source connected to a light guide system with a 700-nm interference filter was used to illuminate the cortical surface through the cranial window. Reflectance images 300 µm below the surface were acquired using a ×2.7 objective and the Imager 3001F (Optical Imaging, Mountainside, NJ) equipped with a 256 × 256 pixels array charge-coupled device (CCD) camera (using VDaq software). The built-in Imager 3001F analysis program (Winmix software) was used to visualize the responses and produce an intrinsic signal image for each whisker by dividing the signal upon stimulation by the baseline signal before stimulation. The intrinsic signal images, and eventually the oriented barrel map outline was superimposed to an image of the vasculature, acquired using a 546-nm interference filter. For Ca^2+^ imaging experiments, this reference image was used to select a field of view with optimal GCaMP and/or ChR2 expression, using a two-photon laser scanning microscope (2PLSM). For behavioral experiments, this reference image was used to determine the stimulation whisker based on the corresponding ChR2-expessing barrel column.

### Slice preparation and whole-cell recordings

#### Somatic recordings

Mice were anesthetized with isoflurane (4% in O_2_; 0.4 L min^−1^) and were transcardially perfused with ice-cold slicing solution containing (in mM): NaCl (83), KCl (2.5) MgSO_4_*7H_2_O (3.3), NaH_2_PO_4_*H_2_O (1), NaHCO_3_ (26.2), CaCl_2_*H_2_O (0.5), glucose (22), sucrose (72), equilibrated with 95% O2/5% CO_2_, pH 7.4. 300 µm-thick coronal brain slices were prepared from 8 to 12 weeks old mice with a vibratome (Leica VT 1000S). Slices were kept for 20 minutes at 37°C in a chamber, and then at room temperature until the time of recording. In the recording chamber, slices were continuously superfused with recording ACSF (32°C) containing (in mM): NaCl (119), KCl (2.5), CaCl_2_ (2.5), MgSO_4_ (1.3), NaH2PO4 (1.0), NaHCO_3_ (26.2), and glucose (22), and equilibrated with 95% O2/5% CO_2_, pH 7.4. Whole-cell recordings were obtained from visually identified NGC-tdT INs in cortical layers 1–3, and PNs in L2/3 and L5b, using an upright microscope (Zeiss Axioskop FS) equipped with differential interference contrast and standard epifluorescence. For current clamp recordings, borosilicate glass patch pipettes had a resistance of 5–6 MΩ when filled with an internal solution containing (in mM): K gluconate (135), KCl (4), HEPES (10), Phosphocreatine (10), Mg-ATP (4), Na-GTP (0.3), and biocytin (2.68). For voltage clamp recordings, borosilicate glass patch pipettes had a resistance of 6–8 MΩ when filled with an internal solution containing (in mM): Cs Methylsulfonate (140), HEPES (10), Phosphocretine (10), MgATP (4), NaGTP (0.3), Biocytin (3), Spermine (0.1). Current clamp recordings were performed upon constant current injection to hold cells at ∼-55 mV. Voltage clamp recordings were performed at −70 mV for excitatory and at 0 mV for inhibitory postsynaptic currents. Optical stimulation was achieved using a 470 nm LED source (Thorlabs) at 1.35 mW/mm^2^ illumination power; pulse duration was 10 ms. Data were acquired using a Multiclamp 700B Amplifier (Molecular Devices), and digitized at 10 kHz (Axon Digidata 1440A), using Clampex 11 acquisition software.

#### Dendritic recordings

Mice were anesthetized with ketamine/xylazine (100 mg kg^-1^/10 mg kg^-1^) mix, and were transcardially perfused with ice-cold slicing solution containing (in mM): KCl (2.8), NaH_2_PO_4_ (1.25), NaHCO_3_ (25), CaCl_2_ (0.5), MgCl_2_ (7), dextrose (7), sucrose (205), ascorbate (103), and sodium pyruvate (3), equilibrated with 95% O2/5% CO_2_, pH 7.4. 250 µm-thick coronal brain slices were prepared from 6 to 8 weeks old mice with a vibratome (Leica VT 1000S), and then moved to a chamber filled with recording ACSF containing (in mM): NaCl (125), KCl (2.5), NaH_2_PO_4_ (1.25), NaHCO_3_ (25), CaCl_2_ (2), MgCl_2_ (1), dextrose (10), and sodium pyruvate (3, equilibrated with 95% O2/5% CO_2_, pH 7.4. Slices were held for 30 minutes at 35°C, and then at room temperature until the time of recording. In the recording chamber, slices were continuously superfused with recording ACSF at 36°C. Whole-cell recordings were obtained from visually identified thick dendritic trunks of L5b PNs, in L2-3, using an upright microscope (Zeiss, Axioskop) equipped with differential interference contrast and standard epifluorescence. Borosilicate glass patch pipettes had a resistance of 15–40 MΩ when filled with an internal solution containing (in mM): K-gluconate (120), KCl (16), HEPES (10), NaCl (8), Phosphocreatine (7), K (2), 0.3 Na-GTP (0.3), 4 Mg-ATP (4). Current clamp recordings were performed upon constant current injection to hold cells at ∼-55 mV. Optical stimulation was achieved using a 475 ± 23 nm coolLED pE-300ultra (CoolLED Ltd); illumination power was 1.6 mW/mm^2^ and pulse duration was 10 ms. Data were acquired using a Multiclamp 700B Amplifier (Molecular Devices), and digitized at 10 kHz (Axon Digidat 1550B), using Clampex 11 acquisition software.

#### Pharmacology

Depending on the experiment, the following drugs were perfused in the bath: CGP 55845 (100 µM; Sigma Aldrich, SML0594), SR 95531 (Gabazine; 10 µM; Sigma Aldrich, 5.05986), APV (50 µM, Sigma Aldrich, A8054), NBQX (10 µM, Abcam, ab120046), TTX (1 µM; Latoxan, L8502), 4-AP (100 µM; Sigma Aldrich, A78403).

#### Quantification of evoked responses

Offline analysis was performed using Clampfit (Version 10.1.0.10, Molecular Devices). Cells were accepted for analysis only if their series resistance was <25 MΩ for somatic or <40 MΩ for dendritic recordings and did not change more than 30% throughout the experiment. Evoked responses were quantified based on the area under the curve (AUC) of postsynaptic potentials or currents within 500 ms after the stimulation onset. For postsynaptic currents, the AUC reflects the total charge (Q) of the response, and was therefore used for computing E-I ratios based on the formula:

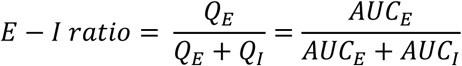

### In vivo Calcium Imaging

We used a custom-built 2PLSM mounted on a modular *in vivo* multiphoton microscopy system (https://www.janelia.org/open-science/mimms-10-2016). The system was equipped with an 8-kHz resonant scanner, a 16× 0.8 NA objective (Nikon, CFI75), and controlled by ScanImage 2016b (http://www.scanimage.org). Excitation was provided by a Ti:Sapphire laser (Chameleon Ultra, Coherent) tuned to 980 nm with an approximate power of 25 mW. Fluorescence signals were collected using GaAsP photomultiplier tubes (10770PB-40, Hamamatsu), with tdTomato and GCaMP signals separated via a dichroic mirror (565dcxr, Chroma) and respective emission filters (ET620/60m and ET525/50m, Chroma). Before imaging, mice were habituated to head restraint under the microscope for 10–15 minutes over 4–5 days.

Tufted dendrites of L5b PNs expressing GCaMP7s were imaged right below the pial surface at approximately 7.5 Hz. The imaging plane was set with a size of 300 × 300 µm (1024 × 512 pixels). L2/3 PNs expressing GCaMP6s were imaged at approximately 30 Hz. The imaging plane was set at 600 × 600 µm (512 × 512 pixels) and positioned at approximately 250 µm below the pial surface. NGC neurons expressing GCaMP6s were imaged at two imaging depths, acquired quasi-simultaneously at approximately 10 Hz using a piezo z-scanner (P-725 PIFOC, Physik Instrumente) for moving the objective over the z-axis in combination with Bessel beam scanning to increase the number of recorded NGC neurons that are inerrantly sparsely distributed. Bessel beam scanning was performed using a plano-convex axicon (1-APX-2NIR-H254-P, Altechna) placed in the beam path, extending the depth of field to around 35 µm. The two planes were set with a size of 600 × 600 µm (512 × 256 pixels) and positioned at approximately 50 and 150 µm below the pia.

Fields of view were recorded for 10 min during which whisker stimulations were delivered every 10 s. For stimulating the whiskers, combs, attached to independent piezo-driven actuators, were placed on both sides of the snout. A stimulation consisted of a train of 5 deflections of 45 ms delivered at 20 Hz.

The stimulation conditions were shuffled across mice. For optogenetic stimulation during imaging, a 470 nm LED (Thorlabs) attached to the detection arm of the microscope was used, and the illumination power was 0.4 mW/mm^2^. During LED illumination, the photomultiplier tubes were closed using mechanical shutters (270 ms blanking period). Fields of view were imaged for 5 min during which 30 stimulations were delivered every 10 s, alternating photostimulation and control conditions 2 times, for a total of 20 min recording time.

### Image processing

Images were processed using custom-written MATLAB scripts and ImageJ (http://rsbweb.nih.gov/ij/). Rigid lateral movement vectors were calculated using the NoRMCorre MATLAB toolbox. Residual bidirectional scanning artifact vectors were calculated using a highest-pixel-line signal correlation between the two scanning directions on the entire frame. Regions of interest (ROIs) were hand-drawn based on shape. The fluorescence time-course of each neuron and dendrite were measured as the average of all pixel values within the ROI. For L2/3 PNs, local neuropil signal was measured for each ROI as the average of pixel values within an automatically defined ring of 15 µm width, 2 µm away from the ROI and excluding overlapping regions with surrounding ROIs. The true GCaMP6s signal of a cell body was then estimated as:

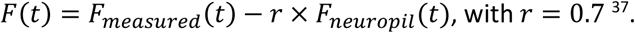

Normalized calcium traces Δ*F*/*F*_0_ were calculated as: (*F*(*t*) − *F*_0_)/*F*_0_, where *F*_0_ is the 30^th^ percentile of the whole *F* trace for L5 tufts dendrites recordings and the average baseline fluorescence signal over 2 s before stimulation for L2/3 pyramidal and NGC neurons. For display, L5 tuft dendrites traces were smoothed using a moving average over 1 s period.

Quantification of evoked responses: Neurons or dendrites whose activity significantly changed across 60 trials were identified using paired t-tests. For NGC neurons, statistical comparisons were performed between the 1 s period before stimulation and either the 1 s or 2 s period following whisker stimulation. A neuron was considered responsive if at least one of these comparisons yielded a significant difference (uncorrected p < 0.05). Neurons that did not show a significant positive or negative change in at least one whisker stimulation condition were classified as non-responding and excluded from further analysis.

For PNs, statistical comparisons were performed between the 1 s period before and the 2 s period following whisker stimulation using a paired t-test (uncorrected p < 0.05). Only neurons exhibiting a significant positive change were included in the analysis. Neurons that did not show a significant increase in any whisker stimulation condition were classified as non-responding and excluded from further analysis, ensuring that calcium imaging analysis was focused on behaviorally relevant PNs activated by whisker stimulation.

Non-responding traces were assigned to cluster 0, indicating no response in the given condition while responding to at least one other condition. Hierarchical clustering was performed on the remaining traces using the average fluorescence responses binned at 20 Hz, applying Euclidean distance and complete linkage to group traces into three clusters for NGCs and L2/3 PNs, and two clusters for L5b tufted dendrites.

Cluster comparison was performed using Fisher’s exact test, revealing that cluster 3 was significantly more prevalent for ipsilateral stimulation, while clusters 2 and 1 were significantly more prevalent for contralateral stimulation. For PNs, the modulation index was calculated as:

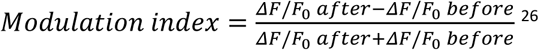

### Histology

For Ca^2+^ imaging and behavior: Mice were anesthetized using isoflurane (4% in O2) for induction and pentobarbital (150 mg/Kg, Esconarkon ad us. Vet. Streuli, Uznach, Switzerland) for maintenance, combined with buprenorphine (0.1 mg/kg, Temgesic, Schering-Plough, NJ, USA) for analgesia. Then they were transcardially perfused with phosphate buffer saline (PBS), followed by paraformaldehyde (PFA; 4% in PBS). After extraction, the brains were post-fixed in the same PFA solution overnight at 4°C, and transferred to PBS. 100 µm-thick coronal slices were cut using a vibratome (Leica VT 1000, Germany). Cell nuclei staining was performed with Hoechst 1:5000 (Invitrogen, MA, USA, #H3570) for 20 min in the dark at room temperature. Samples were imaged using an epifluorescence (Nikon eclipse 80i) and a confocal (Zeiss Confocal LSM800 Airyscan; Axio Imager.Z2 Basis LSM 800) microscope.

For *ex vivo* electrophysiology: Recorded slices were submerged in PFA (4% in PBS) and fixed overnight at 4°C. Biocytin-filled recorded cells were revealed with streptavidin-Alexa 647 conjugate (1:500, Thermo Fisher Scientific) in 0.1% PBS-Tx overnight at room temperature, and cell nuclei staining was performed with Hoechst 1:5000 (Invitrogen, MA, USA, #H3570) for 20 min in the dark at room temperature. Samples were imaged using an epifluorescence (Nikon eclipse 80i) and a confocal (Zeiss Confocal LSM800 Airyscan; Axio Imager.Z2 Basis LSM 800) microscope.

### Behavior

Mice were kept on a reversed light/dark cycle. Habituation of the mice to head restraint began >5 days after the surgery. Head-restrained time at the first day was 5 min and then gradually increased each day until the mice sat calmly for 1 h. Mice were water restricted during subsequent periods of behavioral training. Weight was monitored daily during this period and the amount of water given was adjusted to prevent them from losing more than 15% of their original weight. Altogether, mice received a minimum of 1 ml of water per day corresponding to the amount they drank during the training as rewards plus the amount that the experimenter provided outside of the training sessions. Behavioral control for detecting lick events, delivering whisker stimulation and water reward, and triggering external recording devices, was performed using a data acquisition interface (PCI 6503, National Instruments) and custom-written LabWindows/CVI software (National Instruments). Licks were detected electrically using a metallic plate and spout, creating a 1.2 µA circuit when the tongue touched the spout.

C2 whisker was deflected by displacing a light metal bar (∼3 mg) attached to the whisker, 5 mm away from the snout, using a magnetic coil placed underneath the animal. Local magnetic force was generated by loading the coil with a Gaussian-shaped current pulse (σ = 25 ms) via a custom-designed Arduino-driven high power amplifier (Arduino MKR board). White noise was played during the training sessions to cover potential sound cues generated by the behavioral setup.

For the initial 2 training sessions, the mice received automatic water rewards (∼4 µl for each trial) paired to whisker stimulation with the strongest deflection amplitude (8°, ∼650°/s) to build an association between whisker deflection and reward.

Once the mice establish the association, we introduced a no-lick period and response time window in the behavioral task. After a random inter-trial interval (9.5–13.5 s), a trial started without a preceding cue. Lick events within the 2.5-3.5 s (no-lick period) preceding the scheduled stimulus cancelled the stimulus presentation and reward (time-out). If the mice licked the reward spout within a response window (from 0 to 2 s after the stimulus onset), a drop of water was delivered through the spout. In the following sessions, mice were trained to lick within 2 s after whisker stimulation to trigger the water reward until they reached a hit rate of about 80%. Next, the response window was reduced to 1 s until the mice reached a hit rate of about 80% and cancelled less than 20% of the trials per session. Behavioral sessions were held once or twice a day for each animal.

After the mice learned the task within 1–2 weeks, we tested psychometric functions of the mice by deflecting their C2 whiskers at 7 different amplitudes including no stimulus for catch trials. Whisker stimuli with different intensities were pseudo-randomly interspersed in trials (each intensity presented 3 times in blocks of 21 trials). Typically, the mice needed a couple of sessions before they showed stable psychometric functions. Trials with manipulation (optogenetics or ipsilateral whisker stimulation) were interleaved with control trials.

Psychometric parameters, i.e., detection threshold, false alarm rate and maximum performance, were estimated by fitting detection performance across stimulus intensities with a logistic function ^38^:

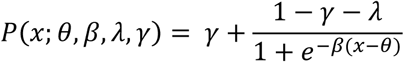

where *x* is stimulus intensity, *P*(*x*) is the detection probability, and θ, β, γ, λ are free parameters that were fitted using a maximum likelihood method. Parameter θ measures the threshold intensity, β measures the slope, γ represents the false alarm rate or chance level, and λ represents the maximum performance of the psychometric function. Curve fitting was performed on results obtained from individual sessions using the maximum likelihood estimation implemented in the Palamedes psychological MATLAB toolbox ^39^.

## Statistical analysis

All statistics were performed using MATLAB and R. For all figures, significance levels were denoted as \**P*<0.05, \*\**P*<0.01, \*\*\**P*<0.001, \*\*\*\**P*<0.0001. No statistical methods were used to estimate sample sizes. Non-parametric tests were used for non-normally distributed data.

## Data availability

The data used to generate the figures is freely available at the CERN data repository Zenodo https://zenodo.org/communities/holtmaat-lab-data/ with 10.5281/zenodo.15074977.

## Code availability

The code that was used for data analysis is freely available at the CERN data repository Zenodo https://zenodo.org/communities/holtmaat-lab-data/ with 10.5281/zenodo.15074977.

## Acknowledgements

We thank Stéphane Pagès, Rashedeh Roshani and Paul Marchand for their help in setting up the Bessel beam imaging. We would like to thank Elodie Husi, Sébastien Pellat and Raphaël Thurnherr for their technical support. This work was supported by the Swiss National Foundation (grants 31003A_173125 and 310030_204562, and NCCR Synapsy 173125/185897).

## Author contributions

Conceptualization, F.M., R.C., A.D. and A.H.; Methodology, F.M., R.C., F.B., and A.H.; Investigation, F.M., R.C., F.B., V.C. and J.P.; Formal Analysis, F.M., R.C., V.C., and J.P.; Resources, A.H.; Writing – Original Draft, F.M., R.C. and A.H.; Writing – Review & Editing, F.M., R.C. and A.H.; Visualization, F.M., R.C. and A.H.; Funding Acquisition, A.H.; Supervision, A.H.

---

**Figure S1.**
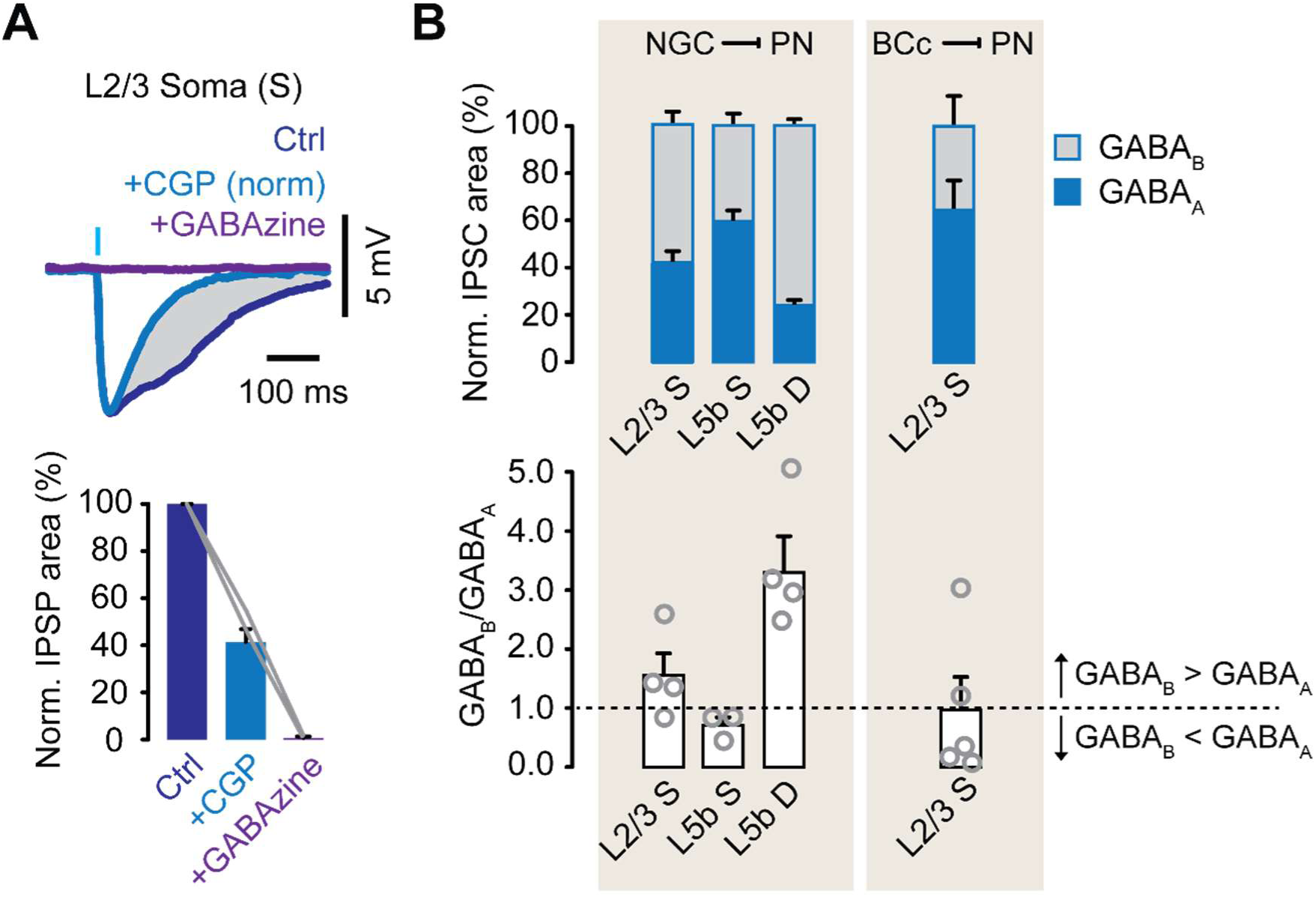
NGC-mediated slow inhibition in pyramidal neurons. (A) Top, Representative example of average IPSPs recorded at membrane potentials of ∼55 mV, upon optical stimulation of L1-3 NGCs before (blue) and after subsequent bath application of CGP-55845 (1 µM; light blue) and GABAzine (10 µM; purple). Bottom, normalized areas of average IPSPs before and after CGP-55845 and GABAzine (*n = 2* cells from *N = 2* mice). (B) Top, comparison of GABA_A_R- and GABA_B_R-dependent components for optical stimulation of NGCs and BCc afferents. Bottom, comparison of GABA_B_/GABA_A_ ratios for optical stimulation of NGCs and BCc afferents.

**Figure S2.**
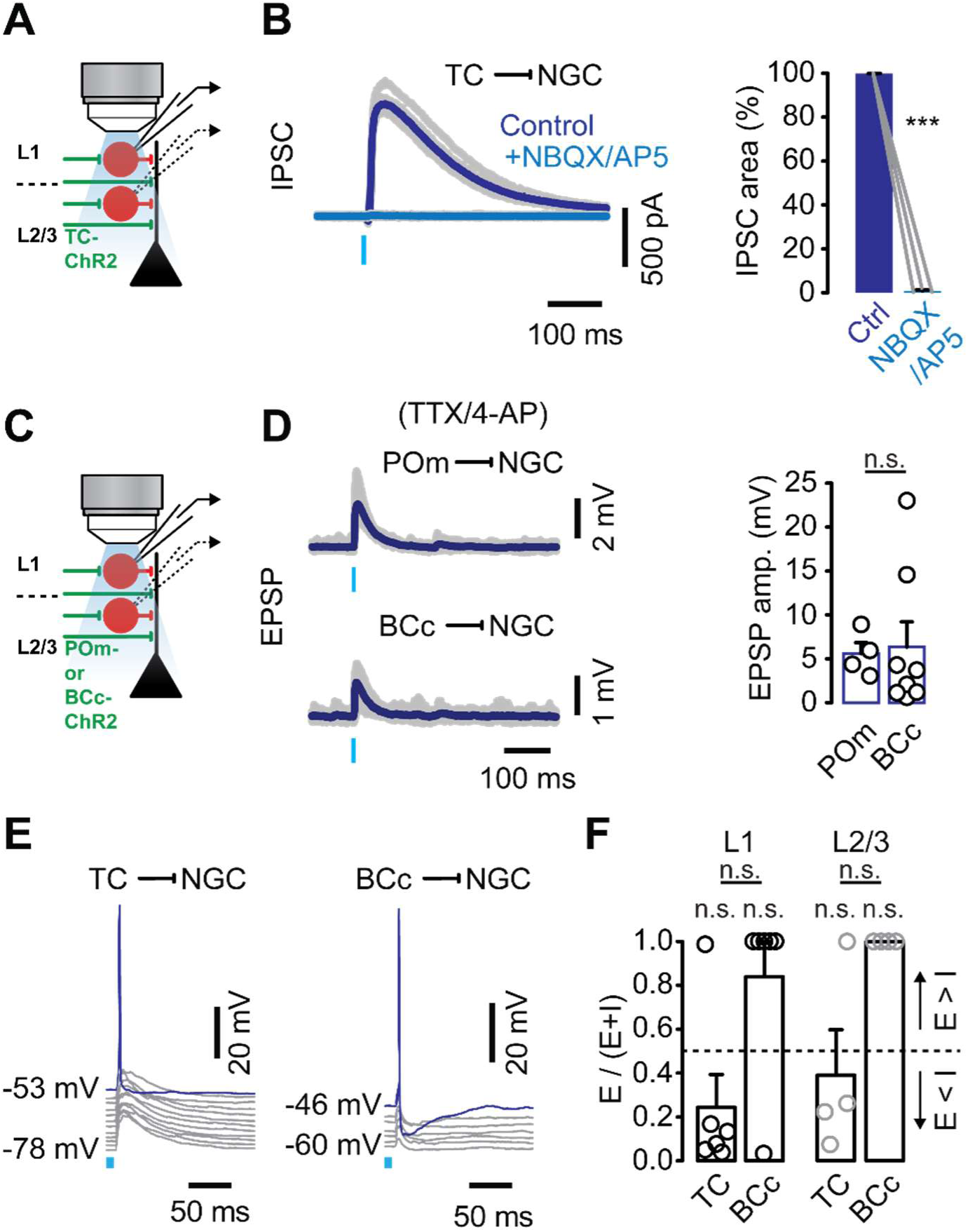
Synaptic inputs from thalamocortical and callosal afferents to L1-3 NGCs. (A) Schematic representation of whole-cell voltage clamp recordings from TdT-expressing L1-3 NGCs. (B) Left, representative example of IPSCs recorded at Vh = 0 mV, upon optical stimulation of TC afferents, before (dark blue) and after bath application of NBQX (10 µM) and AP5 (100 µM) (light blue). Right, average IPSC areas before and after NBQX/AP5 (*n=3* cells from *N=2* mice). (C). Schematic representation of whole-cell current clamp recordings from TdT-expressing L1-3 NGCs in TTX (1 µM) and 4-AP (100 µM). Note that for this experiment we restricted expression of ChR2 to TC afferents from the POm, which constitute the main projections in L1 and were therefore the most likely candidates to monosynaptically contact L1-3 NGCs. (D) Left, representative examples of EPSPs recorded at Vm = ∼55 mV, upon optical stimulation of POm and BCc afferents. Right, average EPSP amplitudes recorded from L1-3 NGCs. (E) Representative examples of spiking activity in L1-3 NGCs upon optical stimulation of TC and BCc afferents, depending on the cells’ resting potential. (F) Comparison between E-I rations of TC and BCc inputs for L1 and L2/3 NGCs (*n = 6* from *N = 3* mice for TC in L1, and *n = 6* from *N = 2* mice for BCc in L1; *n = 4* from *N = 3* mice for TC in L2/3, and *n = 4* from *N = 2* mice for BCc in L2/3). For TC vs BCc in L1, ^n.s.^P = 0.0649, Mann-Whitney test; for TC vs 0.5 and BCc vs 0.5 in L1, ^n.s.^P = 0.4375 and ^n.s.^P = 0.0625, Wilcoxon rank sum test; for TC vs BCc in L2/3, ^n.s.^P = 0.0689, unpaired t-test; for TC vs 0.5 and BCc vs 0.5 in L2/3, ^n.s.^P = 0.8750 and ^n.s.^P = 0.1250, Wilcoxon rank sum test.

**Figure S3.**
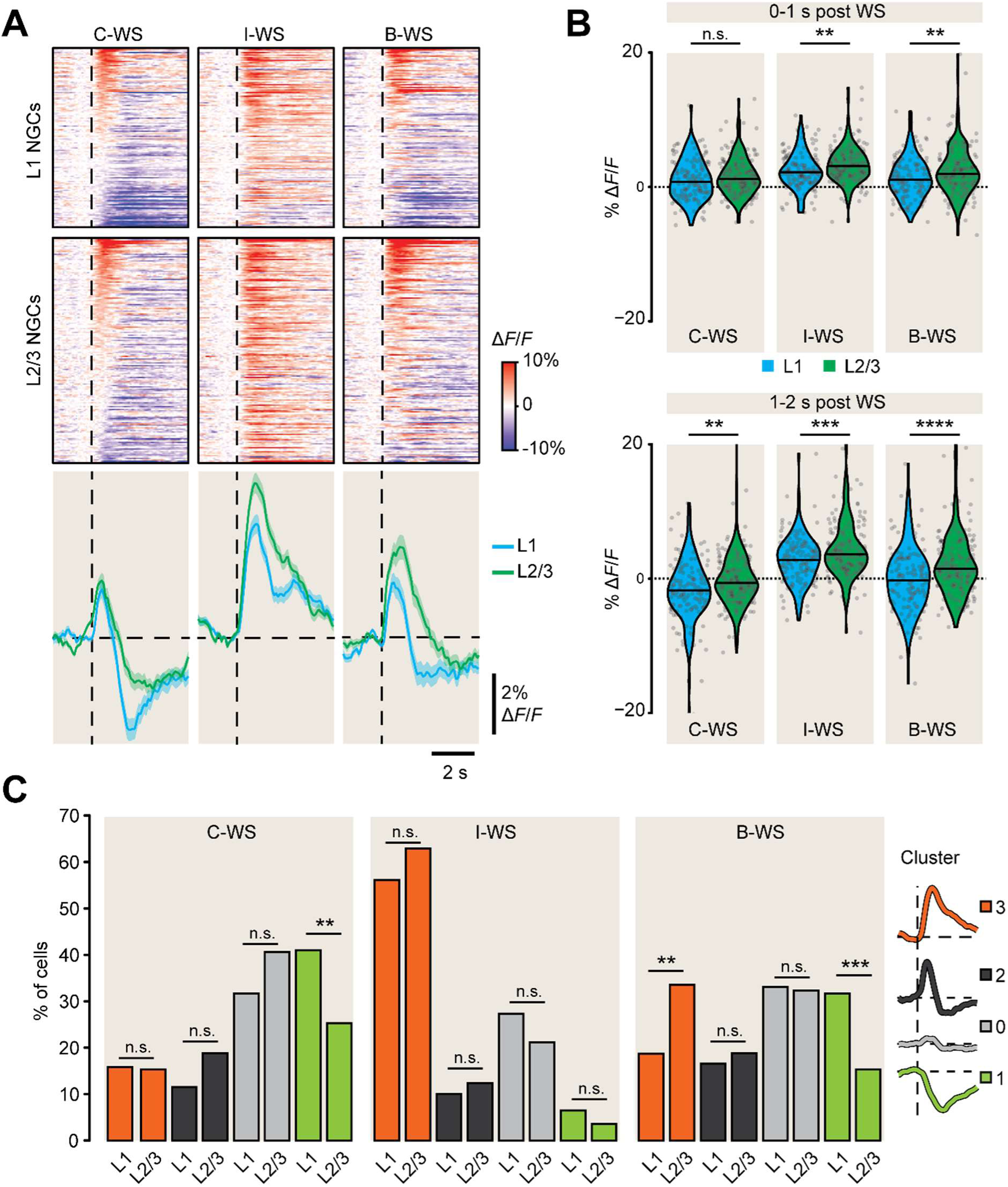
L2/3 NGC activity is less suppressed by contra- and more enhanced by ipsi-lateral whisker stimulation compared to that of L1 NGCs. (A) Top, average calcium responses of each L1 and L2/3 cell to all WS types across 60 trials. Responses are baseline corrected (subtraction of baseline Δ*F/F* −1 to 0 s before stimulus onset) and aligned to stimulus onset (dashed line). Cells are sorted according to their average response 0-1s post C-WS onset. Plots include *n = 139* L1 cells and *n = 170* L2/3 cells from *N=5* mice. Bottom, average response traces of all L1 and L2/3 cells. Shaded areas represent S.E.M. (B) Top, comparison of average responses between L1 and L2/3 NGCs for each WS type, 0-1s post WS onset. Bottom, same for 1-2s post WS onset. C-WS, I-WS, and B-WS: ^n.s.^P = 0.13, **P = 3.1 x 10^-3^, **P = 4.7 x 10^-3^ for 0-1s post WS, and **P = 3.1 x 10^-3^, ***P = 3.4 x 10^-4^, ****P = 6.2 x 10^-5^, for 1-2s post WS; welch two sample t-test. (C) Comparison of cluster proportions between different WS types. Clusters 3,2,0, and 1: ^n.s.^P = 1, ^n.s.^P = 0.08, ^n.s.^P = 0.12, and **P = 4.7 x 10^-3^ for C-WS; ^n.s.^P = 0.24, ^n.s.^P = 0.59, ^n.s.^P = 0.23, and ^n.s.^P = 0.29 for I-WS; **P = 4.4 x 10^-3^, ^n.s.^P = 0.65, ^n.s.^P = 0.90, and ***P = 9.6 x 10^-4^ for B-WS; Fisher’s exact test.

**Figure S4.**
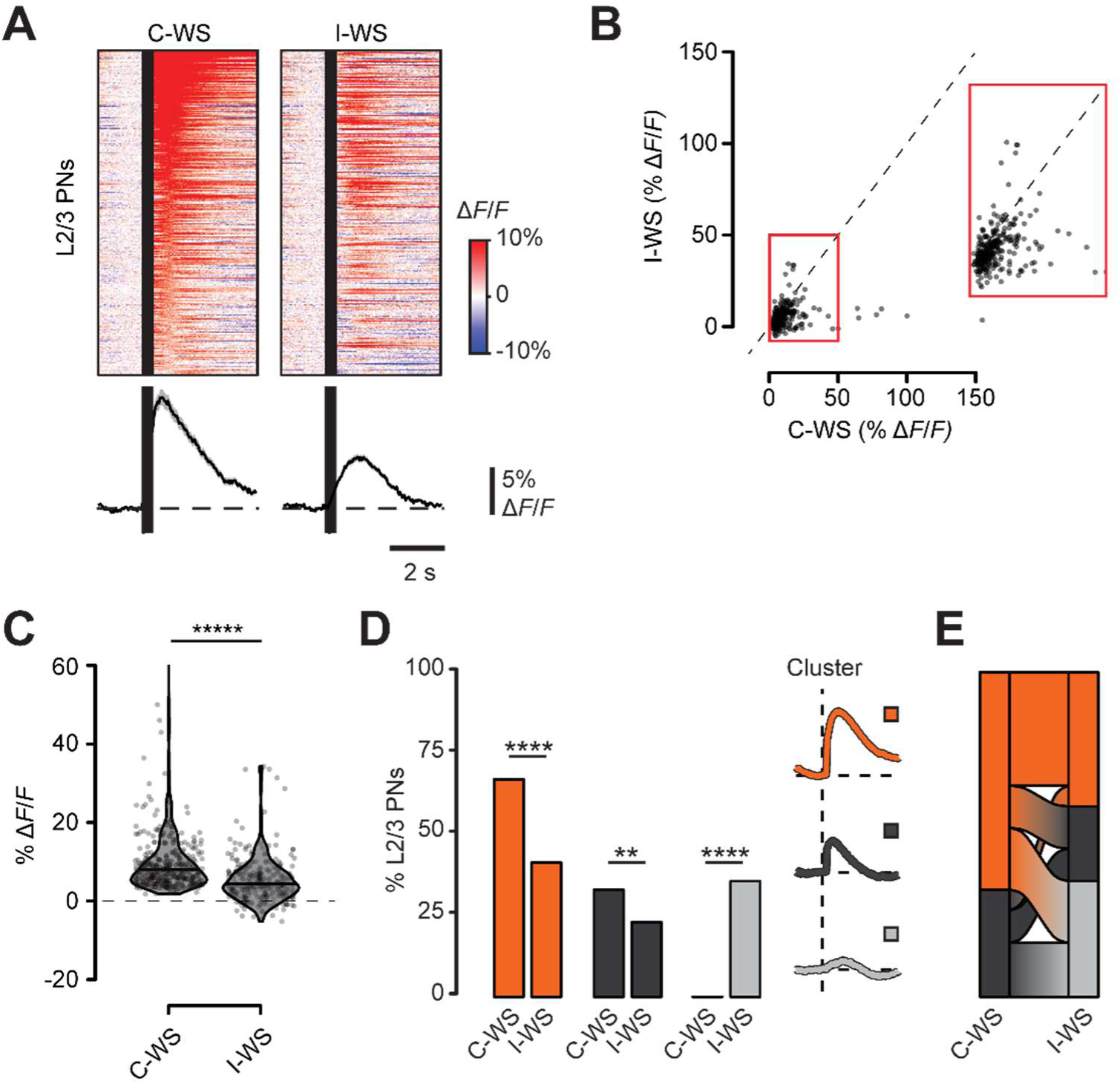
L2/3 PN activity is more enhanced by contra-compared to ipsi-lateral whisker stimulation. (A) Top, average calcium responses of each cell upon C-WS and I-WS across 60 trials. Responses are baseline corrected (subtraction of baseline Δ*F/F* −1 to 0 s before stimulus onset) and aligned to WS onset (black band left border). Black bands indicate periods of no data collection due to the photomultiplier tubes’ shutdown during LED stimulation. Cells are sorted according to their average response 0-2s post C-WS onset. Each plot includes *n = 300* cells from *N = 3* mice. Bottom, average response traces of all cells. Shaded areas represent S.E.M. (B) Scatter plot of average responses 0-2s post WS for C-WS vs I-WS. (C) Comparison of average responses 0-2s post WS onset. ****P = 1.8 x 10^-12^, paired t-test. (D) Comparison of cluster proportions between different C-WS and I-WS. For clusters 2,1, and 0: ****P = 3.8 x 10^-10^, **P = 8.3 x 10^-3^, ****P = 1.15 x 10^-37^, Fisher’s exact test. (E) Alluvial plot illustrating the proportions of cluster changes between C-WS and I-WS. Ribbons representing cluster changes for less than 2% of cells are removed for clarity.

**Figure S5.**
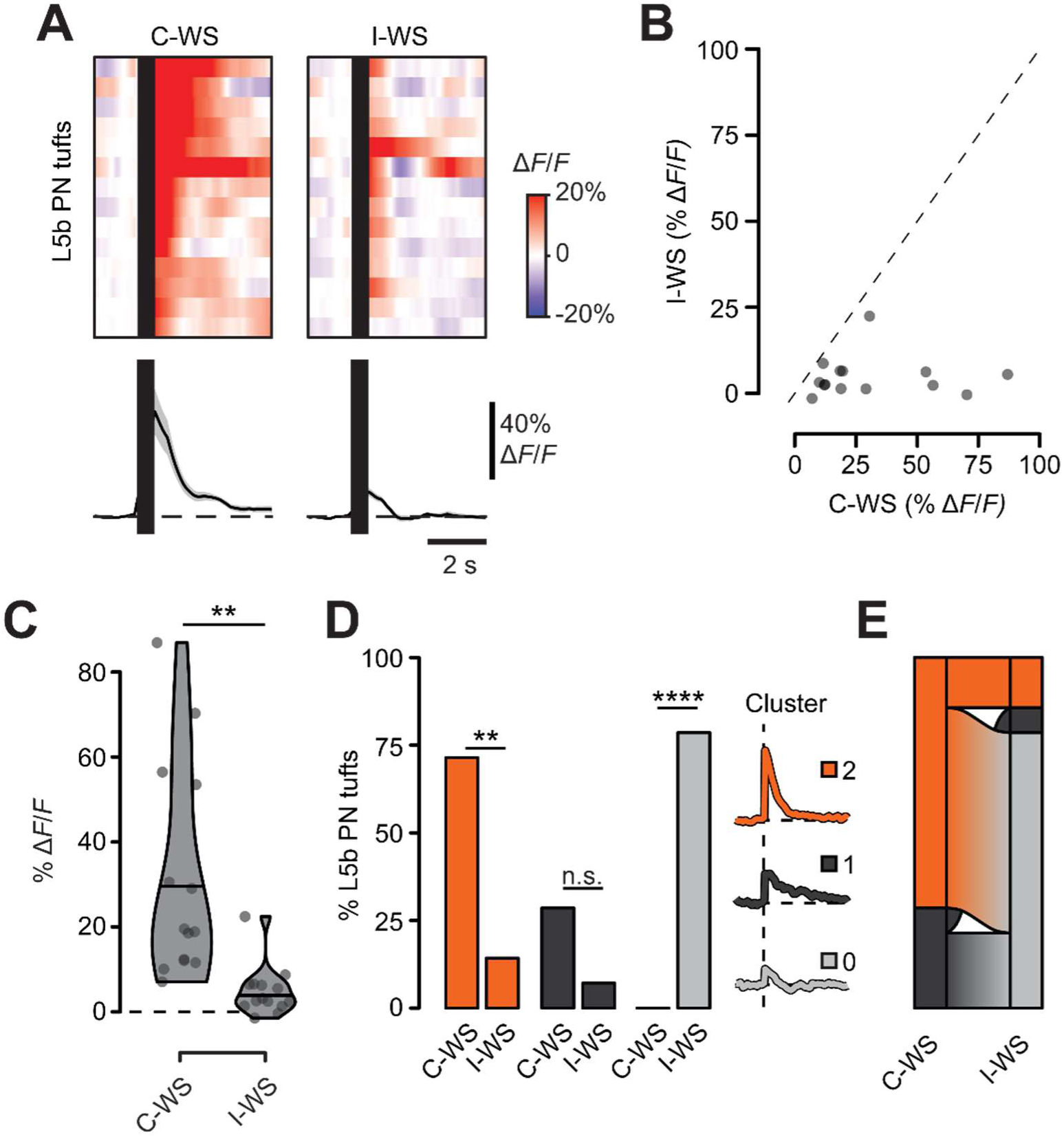
L5b PN tuft activity is more enhanced by contra-compared to ipsi-lateral whisker stimulation. (A) Top, average calcium responses of each tuft to C-WS and I-WS across 60 trials. Responses are baseline corrected (subtraction of baseline Δ*F/F* −1 to 0 s before stimulus onset) and aligned to WS onset (dark band left border). Tufts are sorted according to their average response 0-1s post C-WS onset. Each plot includes *n = 14* tufts from *N = 2* mice. Bottom, average response traces of all tufts. Shaded areas represent S.E.M. (B) Scatter plot of average responses 0-2s post WS for C-WS vs I-WS. (C) Comparison of average responses 0-2s post WS onset. C-WS vs I-WS: **P = 2.2 x 10^-3^, paired t-test. (D) Comparison of cluster proportions between C-WS and I-WS. For clusters 2,1, and 0: **P = 6.3 x 10^-3^, ^n.s.^P = 0.33, ****P = 3.4 x 10^-5^, Fisher’s exact test. (E) Alluvial plot illustrating the proportions of cluster changes between C-WS and I-WS. (C, D) *p < 0.05; **p < 0.01; ***p < 0.001; ****p < 0.0001.

**Figure S6.**
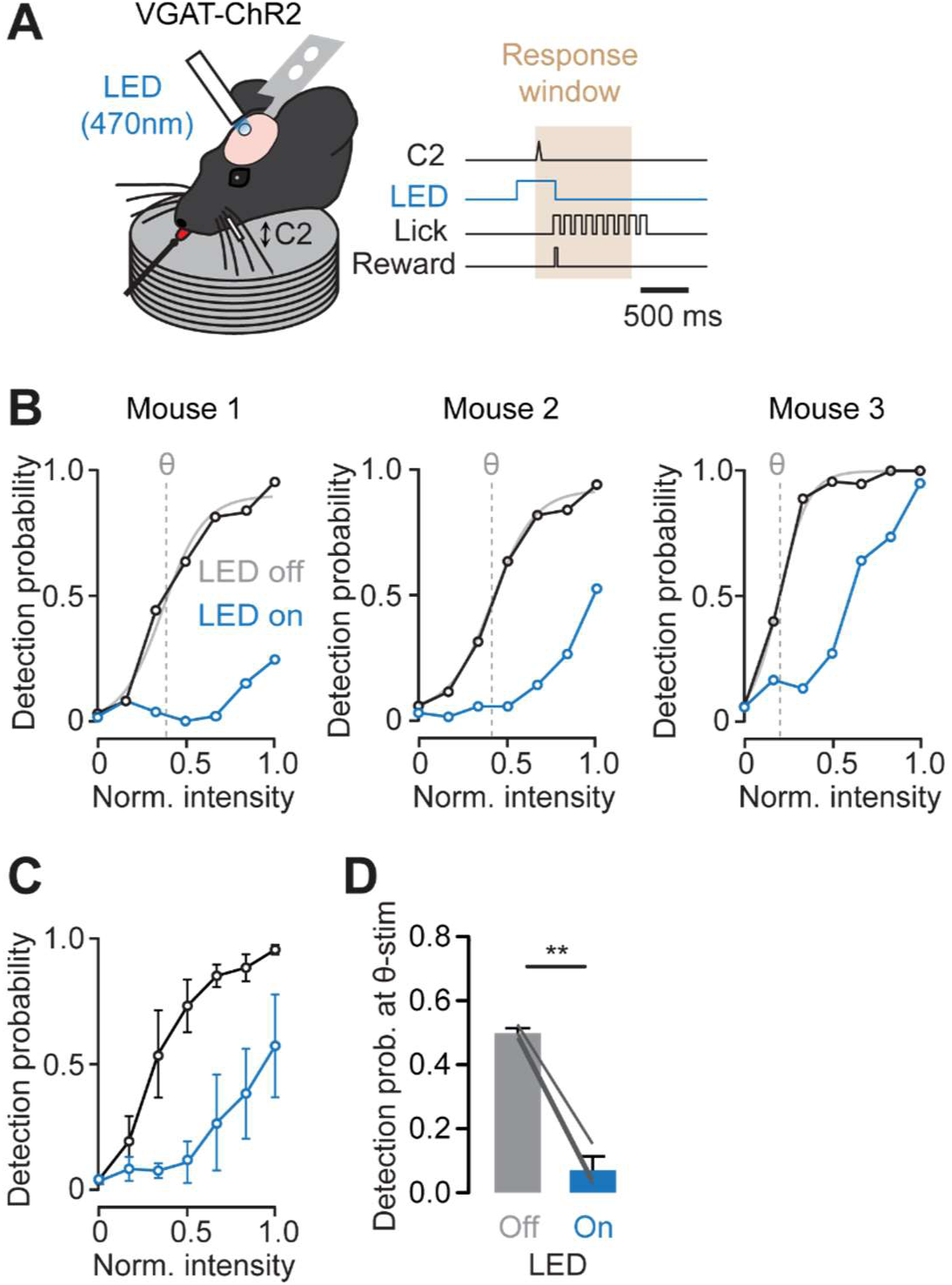
Activity in S1 BC is necessary for whisker-based sensory perception. (A) Schematic representation of the behavioral task combined with optical stimulation of ChR2-expressing VGAT positive interneurons, suppressing the activity in S1 BC. To test the contribution of S1 BC, trials where S1 BC was optically suppressed (LED on) were interleaved with control trials (LED off). (B) Psychometric curves with LED on (blue) and LED off (black) for individual animals. Psychometric functions were computed based on the control psychometric curves (grey lines) to determine the detection threshold (Θ). (C) Average of the psychometric curves with LED on and LED from all animals in (B) (*N* = 3 mice). (D) Detection probability at Θ stimulation determined from the control psychometric curves in (B) showing a significant impairment of sensory perception when S1 BC activity is suppressed. **P = 0.0042, paired t-test.

## References

1. Shuler, M.G., Krupa, D.J., and Nicolelis, M.A.L. (2002). Integration of Bilateral Whisker Stimuli in Rats: Role of the Whisker Barrel Cortices. Cereb. Cortex 12, 86–97. 10.1093/cercor/12.1.86.

2. Pala, A., and Stanley, G.B. (2022). Ipsilateral Stimulus Encoding in Primary and Secondary Somatosensory Cortex of Awake Mice. J. Neurosci. 42, 2701–2715. 10.1523/jneurosci.1417-21.2022.

3. Gauld, O.M., Packer, A.M., Russell, L.E., Dalgleish, H.W.P., Iuga, M., Sacadura, F., Roth, A., Clark, B.A., and Häusser, M. (2024). A latent pool of neurons silenced by sensory-evoked inhibition can be recruited to enhance perception. Neuron 112, 2386–2403.e6. 10.1016/j.neuron.2024.04.015.

4. Cissé, Y., Grenier, F., Timofeev, I., and Steriade, M. (2003). Electrophysiological properties and input-output organization of callosal neurons in cat association cortex. J. Neurophysiol. 89, 1402–1413. 10.1152/jn.0871.2002.

5. Karayannis, T., Huerta-Ocampo, I., and Capogna, M. (2007). GABAergic and pyramidal neurons of deep cortical layers directly receive and differently integrate callosal input. Cereb. Cortex 17, 1213–1226. 10.1093/cercor/bhl035.

6. Petreanu, L., Huber, D., Sobczyk, A., and Svoboda, K. (2007). Channelrhodopsin-2-assisted circuit mapping of long-range callosal projections. Nat. Neurosci. 10, 663–668. 10.1038/nn1891.

7. Palmer, L.M., Schulz, J.M., Murphy, S.C., Ledergerber, D., Murayama, M., and Larkum, M.E. (2012). The Cellular Basis of GABAB-Mediated Interhemispheric Inhibition. Science (80-.). 335, 989–993. 10.1126/science.1217276.

8. Slater, B.J., and Isaacson, J.S. (2020). Interhemispheric callosal projections sharpen frequency tuning and enforce response fidelity in primary auditory cortex. eNeuro 7, 1–11. 10.1523/ENEURO.0256-20.2020.

9. Tamás, G., Lörincz, A., Simon, A., and Szabadics, J. (2003). Identified sources and targets of slow inhibition in the neocortex. Science (80-.). 299, 1902–1905. 10.1126/science.1082053.

10. Abs, E., Poorthuis, R.B., Apelblat, D., Muhammad, K., Pardi, M.B., Enke, L., Kushinsky, D., Pu, D.-L., Eizinger, M.F., Conzelmann, K.-K., et al. (2018). Learning-Related Plasticity in Dendrite-Targeting Layer 1 Interneurons. Neuron 100, 684–699.e6. 10.1016/j.neuron.2018.09.001.

11. Cohen-Kashi Malina, K., Tsivourakis, E., Kushinsky, D., Apelblat, D., Sokoletsky, M., Lampl, I., and Spiegel, I. (2021). NDNF interneurons in layer 1 gain-modulate whole cortical columns according to an animal’s behavioral state. Neuron, 1–28. 10.1016/j.neuron.2021.05.001.

12. Vighagen, R., Gesuita, L., Damilou, A., Banterle, L., and Karayannis, T. (2021). Layer 1 NDNF + Interneurons Control Bilateral Sensory Processing in a Layer-dependent Manner. bioRxiv.

13. Pardi, B., Vogenstahl, J., Dalmay, T., Spanò, T., Pu, D.-L., Bella Naumann, L., Spekeler, H., and Letzkus, J.J. (2020). A top-down thalamo-cortical circuit for associative memory. Science (80-.). 848, 844–848.

14. Ibrahim, L.A., Huang, S., Fernandez-Otero, M., Sherer, M., Qiu, Y., Vemuri, S., Xu, Q., Machold, R., Pouchelon, G., Rudy, B., et al. (2021). Bottom-up inputs are required for establishment of top-down connectivity onto cortical layer 1 neurogliaform cells. Neuron 109, 3473–3485.e5. 10.1016/j.neuron.2021.08.004.

15. Palmer, L.M., Schulz, J.M., and Larkum, M.E. (2013). Layer-specific regulation of cortical neurons by interhemispheric inhibition. Commun. Integr. Biol. 6. 10.4161/cib.23545.

16. Schuman, B., Machold, R.P., Hashikawa, Y., Fuzik, J., Fishell, G.J., and Rudy, B. (2018). Four Unique Interneuron Populations Reside in Neocortical Layer 1. J. Neurosci. 39, 125–139. 10.1523/jneurosci.1613-18.2018.

17. Niquille, M., Limoni, G., Markopoulos, F., Cadilhac, C., Prados, J., Holtmaat, A., and Dayer, A. (2018). Neurogliaform cortical interneurons derive from cells in the preoptic area. Elife 7. 10.7554/eLife.32017.

18. Gelman, D.M., Martini, F.J., Nóbrega-Pereira, S., Pierani, A., Kessaris, N., and Marín, O. (2009). The embryonic preoptic area is a novel source of cortical GABAergic interneurons. J. Neurosci. 29, 9380–9389. 10.1523/JNEUROSCI.0604-09.2009.

19. Fossati, G., Kiss-Bodolay, D., Prados, J., Chéreau, R., Husi, E., Cadilhac, C., Gomez, L., Silva, B.A., Dayer, A., and Holtmaat, A. (2023). Bimodal modulation of L1 interneuron activity in anterior cingulate cortex during fear conditioning. Front. Neural Circuits 17, 1–13. 10.3389/fncir.2023.1138358.

20. Zhang, F., Wang, L.P., Boyden, E.S., and Deisseroth, K. (2006). Channelrhodopsin-2 and optical control of excitable cells. Nat. Methods 3, 785–792. 10.1038/nmeth936.

21. Gambino, F., Pagès, S., Kehayas, V., Baptista, D., Tatti, R., Carleton, A., and Holtmaat, A. (2014). Sensory-evoked LTP driven by dendritic plateau potentials in vivo. Nature 515, 116–119. 10.1038/nature13664.

22. Szabadics, J., Tamás, G., and Soltesz, I. (2007). Different transmitter transients underlie presynaptic cell type specificity of GABAA,slow and GABAA,fast. Proc. Natl. Acad. Sci. U. S. A. 104, 14831–14836. 10.1073/pnas.0707204104.

23. Schulz, J.M., Kay, J.W., Bischofberger, J., and Larkum, M.E. (2021). GABAB Receptor-Mediated Regulation of Dendro-Somatic Synergy in Layer 5 Pyramidal Neurons. Front. Cell. Neurosci. 15, 1–19. 10.3389/fncel.2021.718413.

24. Thériault, G., Thériault, G., De Koninck, Y., De Koninck, Y., and McCarthy, N. (2013). Rapid Volumetric Two-photon Imaging with an Extended Depth of Field. Opt. Life Sci. (2013), Pap. NT1B.6, NT1B.6. 10.1364/NTM.2013.NT1B.6.

25. Towal, R.B., and Hartmann, M.J. (2006). Right-left asymmetries in the whisking behavior of rats anticipate head movements. J. Neurosci. 26, 8838–8846. 10.1523/JNEUROSCI.0581-06.2006.

26. Ayaz, A., Stäuble, A., Hamada, M., Wulf, M.A., Saleem, A.B., and Helmchen, F. (2019). Layer-specific integration of locomotion and sensory information in mouse barrel cortex. Nat. Commun. 10, 1–14. 10.1038/s41467-019-10564-8.

27. Takahashi, N., Oertner, T.G., Hegemann, P., and Larkum, M.E. (2016). Active cortical dendrites modulate perception. Science (80-.). 354. 10.1126/science.aah6066.

28. Takahashi, N., Ebner, C., Sigl-Glöckner, J., Moberg, S., Nierwetberg, S., and Larkum, M.E. (2020). Active dendritic currents gate descending cortical outputs in perception. Nat. Neurosci. 23, 1277–1285. 10.1038/s41593-020-0677-8.

29. Sachidhanandam, S., Sreenivasan, V., Kyriakatos, A., Kremer, Y., and Petersen, C.C.H. (2013). Membrane potential correlates of sensory perception in mouse barrel cortex. Nat. Neurosci. 16, 1671–1677. 10.1038/nn.3532.

30. Shengli, Z., Ting, J.T., Atallah, H.E., Qiu, L., Tan, J., Gloss, B., Augustine, G.J., Deisseroth, K., Luo, M., Graybie, A.M., et al. (2011). Cell-type Specific Optogenetic Mice for Dissecting Neural Circuitry Function. Nat. Methods 8, 745–752. 10.1016/j.bbi.2008.05.010.

31. Babl, S.S., Rummell, B.P., and Sigurdsson, T. (2019). The Spatial Extent of Optogenetic Silencing in Transgenic Mice Expressing Channelrhodopsin in Inhibitory Interneurons. Cell Rep. 29, 1381–1395.e4. 10.1016/j.celrep.2019.09.049.

32. Oryshchuk, A., Sourmpis, C., Weverbergh, J., Asri, R., Esmaeili, V., Modirshanechi, A., Gerstner, W., Petersen, C.C.H., and Crochet, S. (2024). Distributed and specific encoding of sensory, motor, and decision information in the mouse neocortex during goal-directed behavior. Cell Rep. 43, 113618. 10.1016/j.celrep.2023.113618.

33. Hartung, J., Schroeder, A., Péréz, V.R., Poorthuis, R., and Letzkus, J. (2023). Layer 1 NDNF Interneurons are Specialized Top-Down Master Regulators of Cortical Circuits. bioRxiv, 2023.10.02.560136.

34. Groh, A., Bokor, H., Mease, R.A., Plattner, V.M., Hangya, B., Stroh, A., Deschenes, M., and Acsády, L. (2014). Convergence of cortical and sensory driver inputs on single thalamocortical cells. Cereb. Cortex 24, 3167–3179. 10.1093/cercor/bht173.

35. Qi, J., Ye, C., Naskar, S., Inácio, A.R., and Lee, S. (2022). Posteromedial thalamic nucleus activity significantly contributes to perceptual discrimination. PLoS Biol. 20, 1–20. 10.1371/journal.pbio.3001896.

36. Holtmaat, A., Bonhoeffer, T., Chow, D.K., Chuckowree, J., Paola, D., Hofer, S.B., Hübener, M., Keck, T., Knott, G., Lee, A., et al. (2009). Long-term, high-resolution imaging in the mouse neocortex through a chronic cranial window. Nat. Protoc. 4, 1128–1144. 10.1038/nprot.2009.89.Long-term.

37. Chen, T.-W., Wardill, T.J., Sun, Y., Pulver, S.R., Renninger, S.L., Baohan, A., Schreiter, E.R., Kerr, R.A., Orger, M.B., Jayaraman, V., et al. (2013). Ultrasensitive fluorescent proteins for imaging neuronal activity. Nature 499, 295–300. 10.1038/nature12354.

38. Wimmer, R.D., Schmitt, L.I., Davidson, T.J., Nakajima, M., Deisseroth, K., and Halassa, M.M. (2015). Thalamic control of sensory selection in divided attention. Nature 526, 705–709. 10.1038/nature15398.

39. Prins, N., and Kingdom, F.A.A. (2018). Applying the model-comparison approach to test specific research hypotheses in psychophysical research using the Palamedes toolbox. Front. Psychol. 9, 1–14. 10.3389/fpsyg.2018.01250.

